# The protease inhibitor Nirmatrelvir synergizes with inhibitors of GRP78 to suppress SARS-CoV-2 replication

**DOI:** 10.1101/2025.03.09.642200

**Authors:** Doha Al Krad, Kim M. Stegmann, Antje Dickmanns, Claudia Blaurock, Björn-Patrick Mohl, Sina Jasmin Wille, Angele Breithaupt, Tobias Britzke, Anne Balkema-Buschmann, Matthias Dobbelstein

**Author notes:** Corresponding author. Correspondence and requests for materials should be addressed to M. D.

## Abstract

Nirmatrelvir, the active compound of the drug Paxlovid, inhibits the Main protease of SARS-CoV-2 (M^Pro^, 3CL^Pro^, NSP5). Its therapeutic application reduces but does not abolish the progression of COVID-19 in humans. Here we report a strong synergy of Nirmatrelvir with inhibitors of the ER chaperone GRP78 (HSPA5, BiP). Combining Nirmatrelvir with the GRP78-antagonizing drug candidate HA15 strongly inhibits the replication of SARS-CoV-2, to a far greater extent than either drug alone, as observed by diminished cytopathic effect, levels of detectable virus RNA, TCID_50_ titers, accumulation of the non-structural protein 3 (NSP3), as well as Spike and N proteins. The original SARS-CoV-2 strain as well as an Omicron variant were similarly susceptible towards the drug combination. Other GRP78 inhibitors or siRNAs targeting GRP78 also fortified the antiviral effect of Nirmatrelvir. In a hamster model of COVID-19, the combination of Nirmatrelvir with HA15 alleviated pneumonia-induced pulmonary atelectasis more effectively than the single drugs. In conclusion, inhibition of the virus Main protease and cellular GRP78 cooperatively diminishes virus replication and may improve COVID-19 therapy.

## INTRODUCTION

The COVID-19 pandemic has had a profound and lasting impact on global health, causing billions of infections and millions of deaths worldwide. Despite widespread vaccination efforts and the development of antiviral therapies, SARS-CoV-2 continues to circulate, driven by the emergence of novel variants with increased transmissibility and immune evasion. While antivirals have improved the available treatment options, they remain insufficient in fully controlling the disease. Moreover, vaccinations against SARS-CoV-2 are not applicable to certain individuals, e.g. immunocompromised patients. These considerations highlight the ongoing need for effective therapeutic and preventive strategies for current and future pandemics caused by coronaviruses (Chan et al., 2024; Li et al., 2023).

One option to interfere with the SARS-CoV-2 replication cycle (Steiner et al., 2024) is based on the inhibition of a virus protease, thereby precluding the cleavage of viral precursor polyproteins into functional subunits (Ton et al., 2022). SARS-CoV-2 encodes two such proteases; one of them is the Main protease (M^Pro^) or 3C-like protease (3CL^Pro^) or Non-structural protein (NSP) 5, and the second is a papain-like protease (PL^Pro^). Inhibition of the Main protease represents the mechanism of action for the most widely used antiviral small molecule-based drug, Paxlovid. The active compound of Paxlovid, Nirmatrelvir, covalently binds and permanently inactivates M^Pro^ (Owen et al., 2021). Paxlovid also comprises Ritonavir, an inhibitor of the enzyme CYP3A4 that would otherwise degrade Nirmatrelvir. The combination of both molecules confers sufficient protection to significantly reduce the proportion of COVID-19 patients that need hospitalization (Amani and Amani, 2023). However, its application cannot completely avoid disease progression. Moreover, Ritonavir is known to interfere with the turnover of multiple other medications that a patient might require, e.g. antiarrhythmic agents, statins or benzodiazepines, making its clinical application more difficult (Abraham et al., 2022; Prikis and Cameron, 2022). Options for enhancing the efficacy of Nirmatrelvir would therefore be highly desirable, for a stronger antiviral effect and perhaps also to allow the reduction or even elimination of the Ritonavir component of Paxlovid.

The large precursor proteins that encompass the NSPs of SARS-CoV-2 partially localize in the endoplasmic reticulum (ER). NSP4 and NSP6 flank the protease M^Pro^/NSP5 within the precursor protein, and both NSP4 and NSP6 are anchored in the ER via their transmembrane domains (Angelini et al., 2013). On the other hand, the chaperone GRP78 (HSPA5, BiP) assists in the folding of ER-based proteins and supports the transport of proteins into and out of the ER (Ibrahim et al., 2019; Ma and Hendershot, 2004; Zhu and Lee, 2015). This raises the possibility that GRP78 might bind to the NSP precursor proteins, act as chaperone and/or transporter for the intra-ER portions of NSPs, and thus facilitate autocleavage, even when M^Pro^/NSP5 activity is reduced by partial inhibition. In such a situation, pharmacological inhibitors of GRP78 might further prevent the cleavage of NSP precursor proteins and thus interfere with virus replication.

GRP78 has been suggested as a potential target for COVID-19 therapy, and pharmacological agents that lower GRP78 levels can compromise the propagation of SARS-CoV-2 in cell culture models (Ha et al., 2024). However, GRP78 was mostly considered as a co-receptor for SARS-CoV-2 entry. In this concept, GRP78 is exposed on the cell surface and binds to the viral Spike protein, allowing virus entry (Carlos et al., 2021; Han et al., 2022; Han et al., 2023; Shin et al., 2022a; Shin et al., 2021; Shin et al., 2022b). In contrast, the biological function of GRP78 was primarily described as a chaperone within the ER (Ibrahim et al., 2019; Ma and Hendershot, 2004; Zhu and Lee, 2015). It therefore remained to be determined whether the ER-based functions of GRP78 might also contribute to the infectious cycle, and whether the inhibition of this function interferes with virus replication.

A chemical inhibitor of GRP78, HA15, has been described to be effective against malignant pleural mesothelioma (Xu et al., 2019). In this context, it was proposed to enhance proteotoxic stress in tumor cells that produce proteins at a greater rate than non-cancerous cells. In general, the inhibition of chaperones, or heat shock proteins, represents a viable strategy for cancer treatment (Dobbelstein and Moll, 2014). However, this does not preclude the use of cancer drugs for treating infectious diseases, as we have previously described for folate antagonists or DHODH inhibitors (Schrell et al., 2025; Stegmann et al., 2021; Stegmann et al., 2022; Zibat et al., 2023).

Here we demonstrate a strong synergistic interference of Nirmatrelvir and HA15 with the replication of SARS-CoV-2. The drugs cooperate in diminishing the accumulation of virus proteins, virus RNA and virus progeny, and they reduce COVID-19-like lung pathology in an animal model, raising the perspective of their clinical use against SARS-CoV-2 infections.

## RESULTS

### Nirmatrelvir and the GRP78 inhibitor HA15 synergistically interfere with SARS-CoV-2 replication

We tested the efficacy of Nirmatrelvir and the GRP78 inhibitor HA15, alone or together, on the Cytopathic Effect (CPE; Fig. 1A) as well as the release of virus RNA (Figs. 1B; S1A, S1B) upon infection of Vero E6 (Vero C1008, Vero 76, clone E6) cells with a Wuhan-like strain of SARS-CoV-2. At concentrations up to 0.5 µM (Nirmatrelvir) and 5 µM (HA15), neither of the drugs eliminated the CPE, nor did it diminish the release of virus RNA more than 10-fold. In contrast, the combination of both drugs virtually eliminated the CPE, and the amount of released virus RNA was reduced more than 1000-fold. Synergy was confirmed by the Bliss independence model (Fig. 1C). Comparable degrees of synergistic antiviral efficacy were also found when assessing the release of infectious particles by TCID_50_ assays (Fig. 1D). The drugs did not reveal detectable cytotoxicity at these concentrations, as revealed by the quantification of released lactate dehydrogenase (LDH; Fig. 1E). Similar suppression of virus RNA yield was observed in the cell line Calu-3 (Figs. S2A-C; S3), derived from human lung cancer (Kreft et al., 2015). We conclude that inhibition of the virus M^Pro^ as well as GRP78 synergistically interferes with the replication of SARS-CoV-2.

**Figure 1:**
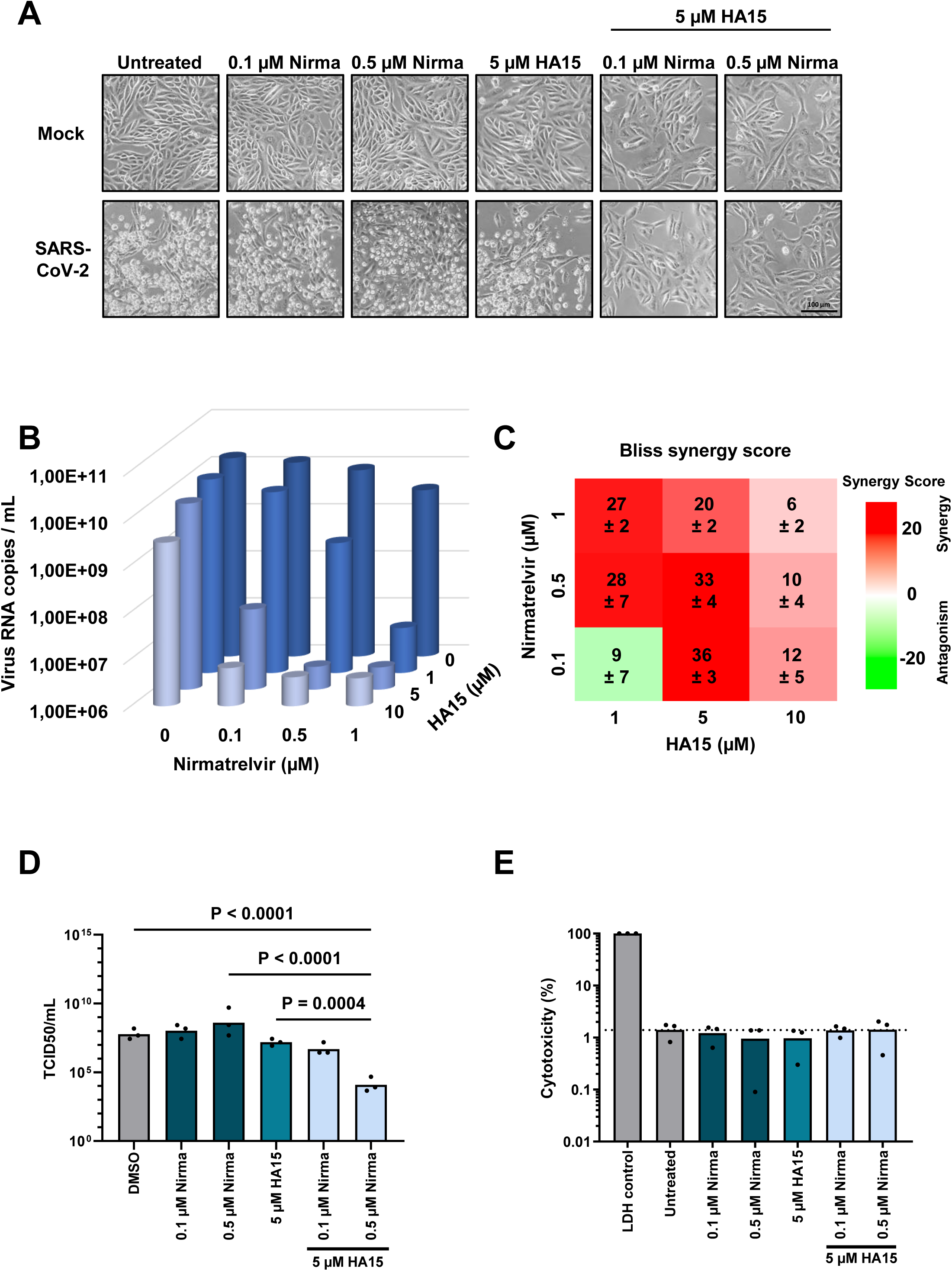
Synergistic interference of Nirmatrelvir and the GRP78 inhibitor HA15 with SARS-CoV-2 replication. A. Diminished Cytopathic Effect (CPE) in infected cells upon simultaneous treatment with Nirmatrelvir and HA15. Vero E6 cells were treated with both drugs, alone or in combination, at the indicated concentrations. The cells were then infected with SARS-CoV-2, Göttingen strain GOE_001 (Stegmann et al., 2021), at MOI 0.02. 48 h post infection (hpi), cell morphology was assessed by phase contrast microscopy. Scale bar, 100 µM. B. Strongly reduced yield of virus RNA in response to simultaneous treatment. Cells were infected and 2 hours later treated with the indicated drug concentrations. At 48 hpi, the supernatant was collected. Virus RNA was quantified by RT-PCR (mean, n = 3). For single data points and P values, cf. Fig. S1A, S1B. Similar effects were observed in Calu-3 cells (Fig. S2A-C; S3). C. The Bliss independence model was used to calculate the synergy score. Bliss values > 10 indicate strong drug synergism (mean ± SEM). D. Synergistic reduction of virus yield by Nirmatrelvir and HA15. Upon Vero E6 cell treatment and infection as in A, infectious virus units in the supernatants were determined by assessing the TCID_50_. E. Lack of observable cytotoxicity. Vero E6 cells were treated with Nirmatrelvir (Nirma) and/or HA15, followed by quantifying Lactate dehydrogenase (LDH) in the supernatant, as a readout for cell lysis. For comparison, fully lysed cells were subjected to the analysis in parallel (control LDH).

### Nirmatrelvir and HA15 cooperatively reduce virus protein synthesis

To confirm the combined antiviral activity of Nirmatrelvir and HA15, we treated and infected Vero cells, followed by immunofluorescence (Fig. 2A) and immunoblot (Fig. 2B) analyses. Both the virus Spike protein and the Nucleoprotein were suppressed below detectable levels by the drug combination, at concentrations that did not diminish virus protein levels when used alone. These findings provide additional evidence that inhibitors of M^Pro^ and GRP78 cooperate to suppress virus replication.

**Figure 2:**
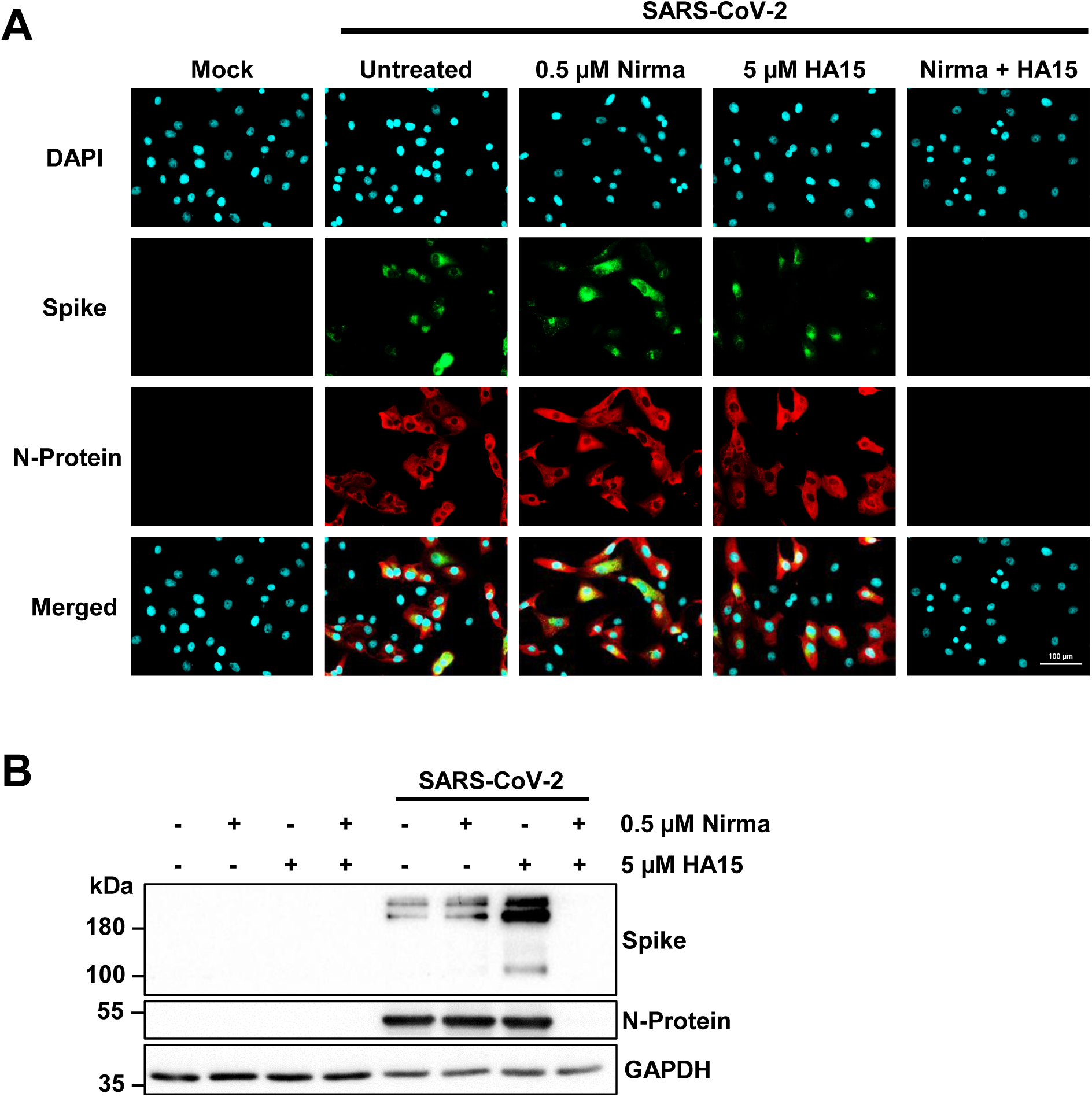
Reduced virus protein synthesis in response to Nirmatrelvir in combination with HA15. A. Elimination of detectable virus protein by the drug combination, as determined by immunofluorescence analysis. Vero E6 cells were treated with the drugs and infected as in Fig. 1A for 48 h. The cells were fixed and stained by antibodies to detect the Spike protein (S) and the Nucleoprotein (N) of SARS-CoV-2. Scale bar, 100 µM. B. Strongly reduced virus protein levels upon combined drug treatment, assessed by immunoblot analysis. Upon treatment and infection as in Fig. 1A, cell lysates were subjected to SDS-PAGE and blotting, and the N and S proteins of SARS-CoV-2 were detected.

### Nirmatrelvir and HA15 diminish the replication of the SARS-CoV-2 variant Omicron

Novel SARS-CoV-2 Variants of Concern (VOC) such as the Omicron variants have replaced the original Wuhan-like strains during the pandemic. With the perspective of clinical use, we tested whether the combination of Nirmatrelvir and HA15 also interferes with Omicron B.1.1.529 replication. Assessing the released virus RNA revealed similar drug synergy as with the original SARS-CoV-2 strain (Figs. 3A, 3B; S4A, S4B).

**Figure 3:**
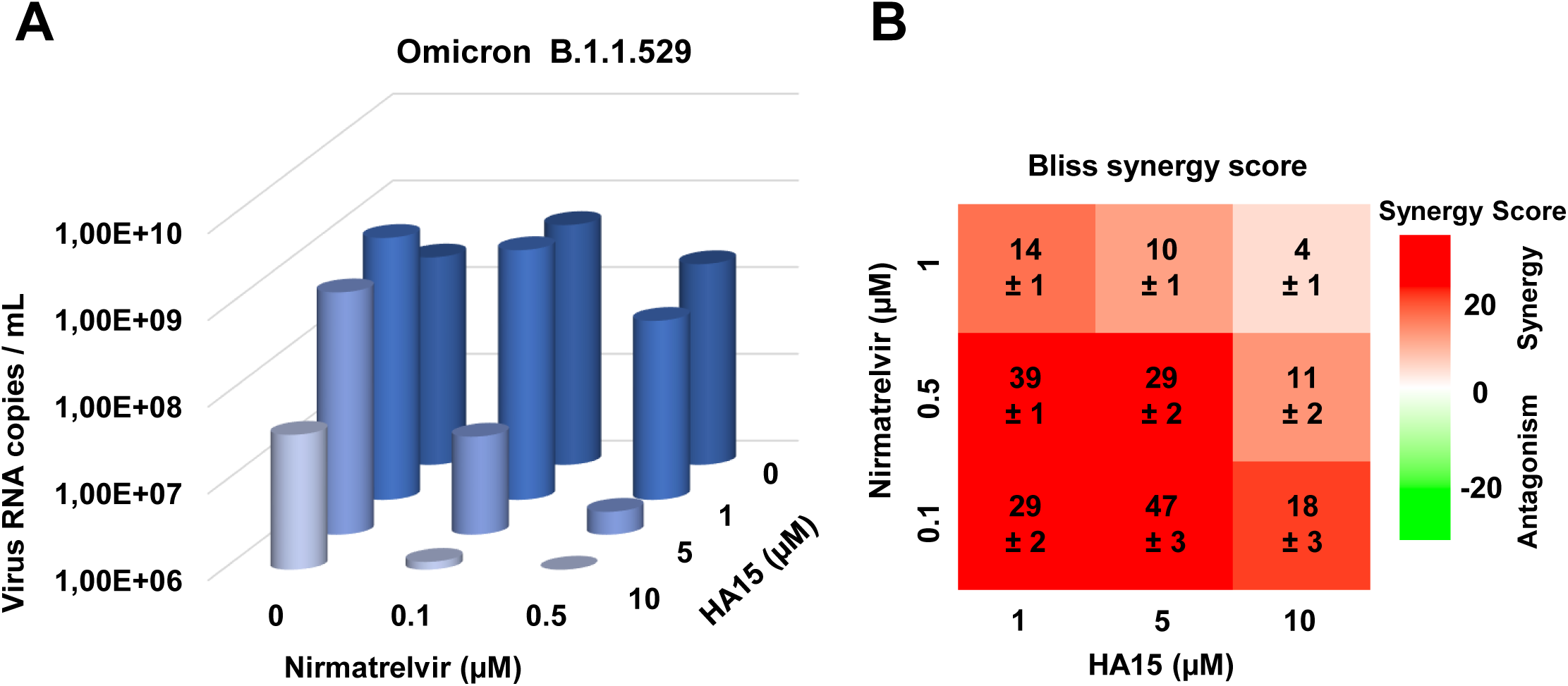
Synergy of Nirmatrelvir and HA15 to counteract the replication of the SARS-CoV-2 variant Omicron. A. Vero E6 cells were infected with the SARS-CoV-2 Omicron variant B.1.1.529 for 2 h. Afterwards, the inoculum was removed, and the cells were washed with PBS and incubated with Nirmatrelvir and/or HA15 for 48 h. Supernatants were then collected, and viral RNA was quantified (mean, n = 3). Single data and P values are provided in Fig. S4A, S4B. B. The Bliss independence model was used to calculate the synergy score. Bliss values > 10 show strong drug synergism (mean ± SEM).

### Targeting GRP78 by additional drugs or siRNA fortifies the antiviral efficacy of M^Pro^ inhibitors

Next, we sought to ensure that the enhancement of Nirmatrelvir efficacy by HA15 is a true on-target effect, and that it can be ascribed to diminished GRP78 function. To test this, we first used an alternative inhibitor of GRP78, Celastrol (Luo et al., 2022), with no obvious chemical similarity to HA15. Indeed, this inhibitor was also capable of synergizing with Nirmatrelvir in antiviral efficacy (Figs. 4A; S5A, S5B). This strongly argues that the drug synergy is actually due to the inhibition of GRP78 and M^Pro^. To further support this observation, we depleted GRP78 from Vero cells by siRNA transfection, followed by infection in the presence or absence of Nirmatrelvir (Figs. 4B, 4C; S5C, S5D). We observed that GRP78 depletion enhanced the antiviral efficacy of Nirmatrelvir, again arguing that the HA15-target GRP78 is a key supporter of virus replication when interfering with M^Pro^ activity. On the other hand, HA15 not only synergized with Nirmatrelvir, but also with another pharmacological inhibitor of M^Pro^, 11a (Dai et al., 2020) (Figs. 4D; S6A, S6B). Taken together, the inhibition of GRP78 and M^Pro^ gives rise to antiviral synergy, regardless of the particular choice of inhibitors.

**Figure 4:**
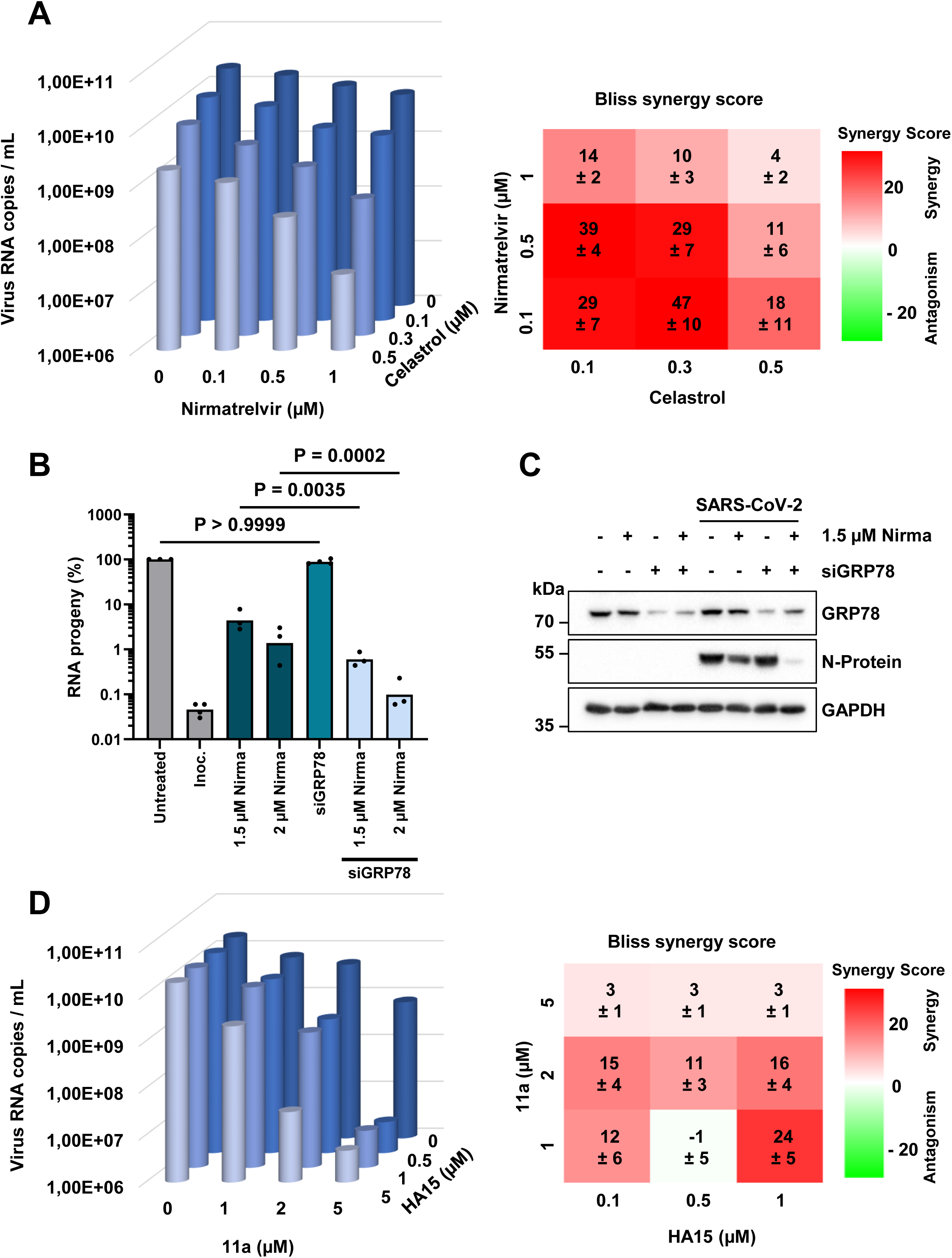
Enhanced antiviral efficacy of M^Pro^ inhibitors, by GRP78-inhibitors and siRNAs targeting GRP78. A. Cooperation of Nirmatrelvir with the GRP78 inhibitor Celastrol. Vero E6 cells were treated and infected as in Fig. 1A but replacing HA15 with Celastrol. The release of virus RNA was quantified (mean, n = 3). The Bliss independence model was used to calculate the synergy score. Bliss values > 10 show strong drug synergism (mean ± SEM). Single data and P values are shown in Fig. S5A, S5B. B. Enhanced Nirmatrelvir efficacy by GRP78-knockdown. Vero E6 cells were transfected with siRNAs to deplete GRP78. 24 h later, the cells were treated with Nirmatrelvir at the indicated concentrations. 8 h later, the cells were infected at MOI 0.02. At 48 hpi, virus RNA in the supernatant was quantified. For P values, cf. Fig. S5C. C. Efficacy of the GRP78 knockdown, determined by immunoblot analysis. A replicate with higher Nirmatrelvir concentration is provided in Fig. S5D. D. Synergistic antiviral effect of HA15 with the M^Pro^ inhibitor 11a. Addition of inhibitors and infection of Vero E6 cells was done as indicated and following the protocol as in Fig. 1A (mean n = 3). Bliss synergy scores were calculated (mean ± SEM). Single data and P values are provided in Fig. S6A, S6B.

### HA15 effectively diminishes virus replication after virus entry

Previously, GRP78 was described as a potential co-receptor for SARS-CoV-2 on the cell surface (Carlos et al., 2021; Han et al., 2022; Han et al., 2023; Shin et al., 2022a; Shin et al., 2021; Shin et al., 2022b). On the other hand, GRP78 was initially reported to serve as a chaperone within the ER (Ma and Hendershot, 2004; Zhu and Lee, 2015). Our next goal was therefore to investigate whether the intracellular functions of GRP78 might contribute to the infectious cycle, and whether blocking these functions could interfere with virus replication. We first inoculated Vero E6 cells with the virus, and only afterwards added HA15 and/or Nirmatrelvir. We reasoned that this should circumvent any impact of the drug on a putative receptor function. The virus RNA was harvested at 6 or 8 hours post-infection (hpi), excluding multiple rounds of virus replication. Remarkably, the GRP78 inhibitor HA15 still synergized with Nirmatrelvir to block the infection cycle, even after permitting virus entry (Fig. 5A-E). Moreover, the drug combination suppressed intracellular RNA levels far more intensely at 8 hpi than at 6 hpi (Figs. 5C, 5E; S7A, S7B). Taken together, this strongly argues in favour of an intracellular function of GRP78 during virus replication.

**Figure 5:**
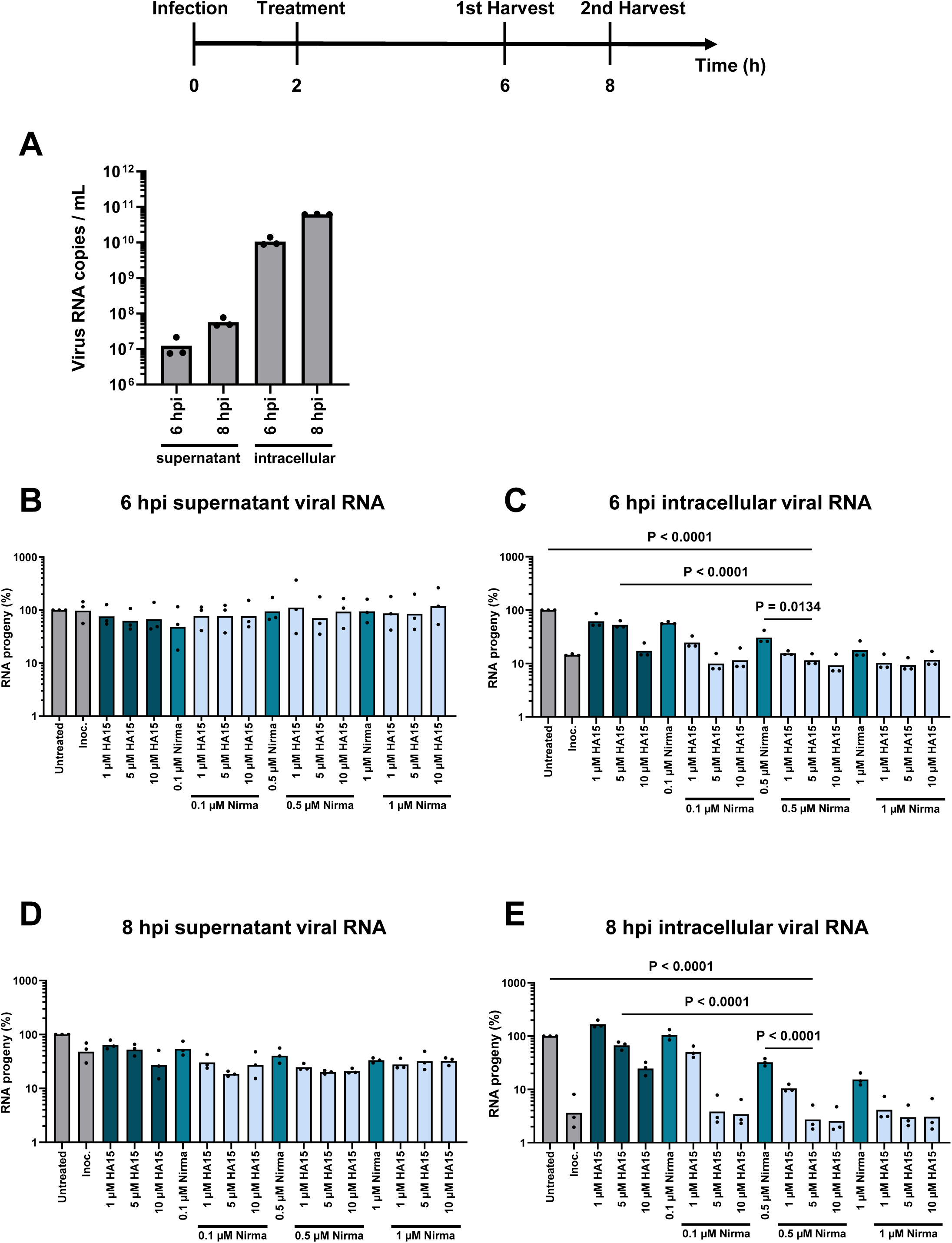
Cooperation with Nirmatrelvir when applying HA15 after virus entry. A. Upon infection of Vero E6 cells with SARS-CoV-2 at MOI 2, without drug treatment, the amount of virus RNA was determined at 6 hpi and 8 hpi, in the supernatant and in the cell lysate, for a baseline to validate the subsequent experiments. B-E. Vero E6 cells were infected with SARS-CoV-2 at MOI 2. At 2 hpi, the virus inoculum was removed, and the drugs were added to the cells as indicated. At 6 hpi (B, C) and 8 hpi (D, E), the viral RNA released to the supernatant (B, D) and the intracellular viral RNA (C, E) were quantified. Of note, the impact of Nirmatrelvir and HA15 on the intracellular virus RNA levels strongly increased from 6 hpi to 8 hpi, arguing that the drugs mainly act on replication steps after virus entry. P values for panel C and E are provided in Fig. S7A, S7B.

### Nirmatrelvir and HA15 together diminish the accumulation of virus proteins

To elucidate the mechanism underlying the synergistic effect of Nirmatrelvir and HA15, we followed viral protein synthesis during the early stages of the replication cycle. As shown in Fig. 2, the inhibition of M^Pro^ and GRP78 reduced the accumulation of Spike and Nucleoprotein, both structural proteins. To further explore the impact of these drugs on a non-structural protein (NSP), we analysed NSP3 levels during the early replication stages. Notably, the combination of Nirmatrelvir and HA15 reduced the NSP3 signal as early as 6 hpi, as revealed by immunofluorescence (Figs. S8; S9) as well as by immunoblot analysis (Figs. 6; S10). In contrast, Nucleoprotein levels were only compromised by the drug combination from 8 hpi onwards (Fig. 6). These results are at least compatible with the view that inhibition of M^Pro^ and GRP78 diminishes NSP3 accumulation early during infection, perhaps by interfering with the processing of NSPs.

**Figure 6:**
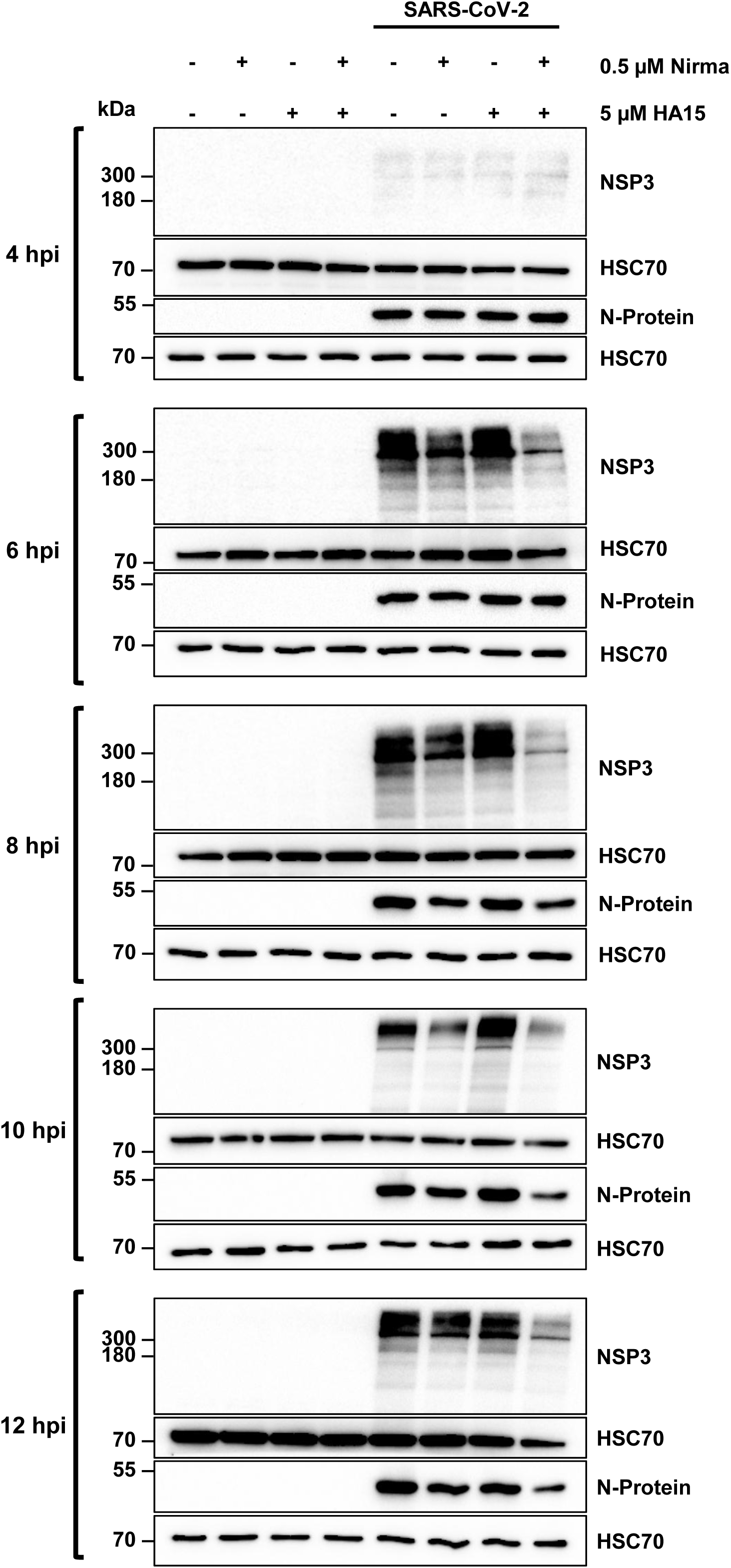
Diminished accumulation of cleaved NSP3 in the presence of Nirmatrelvir and HA15. Vero E6 cells were infected with SARS-CoV-2, GOE_001, at MOI 2. At 2 hpi, the inoculum was removed, and the cells were washed with PBS. Subsequently, the cells were treated with 0.5 µM Nirmatrelvir and 5 µM HA15, either individually or in combination. To investigate the effect of the treatment during the early stages of the viral replication cycle, cells were harvested at 4, 6, 8, 10, and 12 hpi, and protein samples were analysed by immunoblotting. NSP3 was detected at its expected molecular weight of approximately 200 kDa. When both drugs were combined, this signal was diminished as early as 6 hpi, whereas the amount of detectable N-protein was still unchanged at this time. This suggests that the inhibition of M^Pro^ and GRP78 diminishes the accumulation of the SARS-CoV-2 NSP3 early during infection. Cf. Figs. S8; S9; S10 for a similar experiment.

### Nirmatrelvir and HA15 cooperate for therapeutic efficacy in a hamster model of COVID-19

Finally, we evaluated the combination of Nirmatrelvir and the GRP78 inhibitor HA15 in an animal model of COVID-19. Golden Syrian hamsters were pre-treated with Nirmatrelvir, HA15, and/or Ritonavir, an inhibitor of Cytochrome P450 CYP3A4 that prolongs the half-life of Nirmatrelvir (Loos et al., 2022), for one day. Then, the animals were infected with SARS-CoV-2, followed by a daily treatment (Nirmatrelvir twice per day; HA15 and Ritonavir once daily) until the end of the study at day 6 post infection (p.i.). Within the limits of the animal cohort sizes, we did not observe beneficial effects of the HA15 – as monotherapy or when combined with Nirmatrelvir and/or Ritonavir – on animal weight or anti-SARS-CoV-2 serology (Fig. S11A, S11B), nor did we find significant differences in virus titers from nasal swabs or post mortem organ isolates (Fig. S12). However, differences were observed in lung pathology. At day 6 p.i., the lungs of the animals were subjected to histopathological examination. While none of the treatments fully prevented pneumonia-related atelectasis, the extent differed depending on the treatment, and the lungs of the group treated with the combination of all three drugs had lower atelectasis scores compared to the others, with the highest level of significance compared to non-treated animals (Figs. 7A, 7B; S13A; Suppl. Table 1). Inflammatory SARS-CoV-2-typical lesions were observed in all infected groups, regardless of treatment (Fig. S13B). These lesions were characterized by alveolar immune cell infiltrates and alveolar edema (Fig. S13C), perivascular immune cell infiltrates, vasculitis and immune cell rolling (Fig. S13D) as well as peribronchial immune cell infiltrates (Fig. S13E). Hyperplasia and hypertrophy of type 2 pneumocytes (Fig. S13F) and bronchial epithelium were signs of regeneration. In sum, while HA15 alone did not detectably affect the detected virus titers (within the limitations of animal cohorts and large overall variations), it did ameliorate lung pathology when combined with Nirmatrelvir and Ritonavir. This argues in favor of further evaluating combinations of M^Pro^ inhibitors with GRP78 inhibitors, with the perspective of future clinical use.

**Figure 7:**
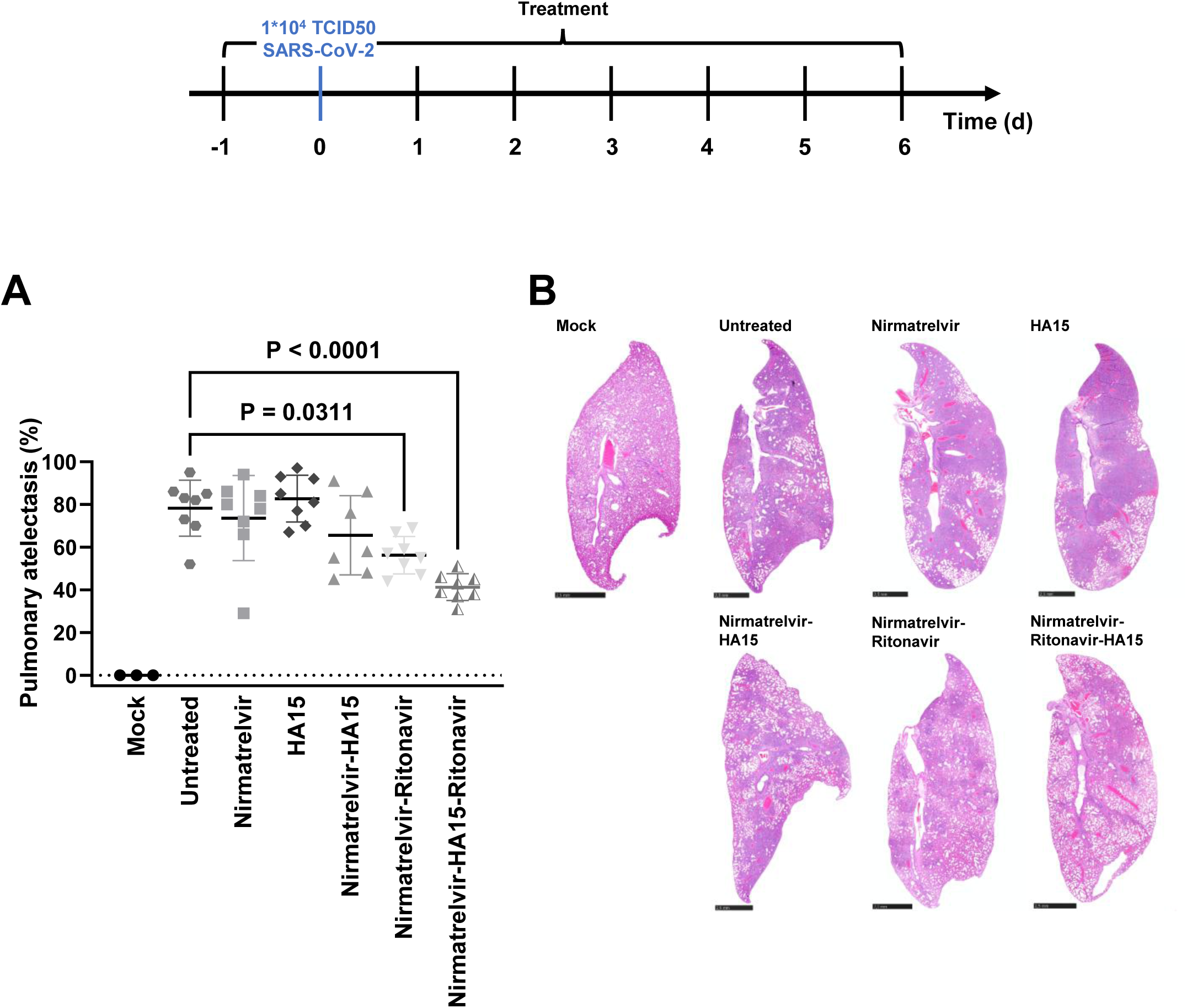
Therapeutic effect of Nirmatrelvir and HA15 in a hamster model of COVID-19. Golden Syrian hamsters were treated with Nirmatrelvir (twice daily – orally – 250 mg/kg), Ritonavir (once daily –orally – 50 mg/kg) and/or HA15 (once daily – intraperitoneally – 100 mg/kg) in various combinations at -1 dpi. 24 h later, the animals were infected with SARS-CoV-2 (Wuhan) and then continuously treated with the drugs until 6 dpi. At this point, the lung histopathology was determined. Additional data on body weight, serology and virus titers are provided in Figs. S11A, S11B; S12. A. Pneumonia-induced pulmonary atelectasis indicated as % affected area. Graphs indicate the arithmetic mean and standard deviation in each case. One-way ANOVA test with Tukey’s multiple comparisons was performed to identify significant difference between triple-treated animals and the untreated or monotherapy groups. Single data and P values are provided in Fig. S13A-F and Suppl. Table 1. B. Histopathology, lung whole slide images showing atelectasis to different extents in all animals except the non-infected mock animals, hematoxylin-eosin stain, bars 2.5 mm.

## DISCUSSION

Our results indicate that inhibitors of the SARS-CoV-2 Main protease M^Pro^/NSP5 and the ER-chaperone GRP78 strongly synergize to interfere with virus replication in cell culture. They also alleviate COVID-19-associated pneumonia in an animal model. This raises the perspective of using such combinations of inhibitors for treating COVID-19.

A role for GRP78 in SARS-CoV-2 infection was previously described (Shin et al., 2022a; Shin et al., 2022b). However, its function in the virus replication cycle was ascribed to facilitating virus entry (Carlos et al., 2021), rather than virus protein synthesis. According to the previous studies, GRP78 acts as a co-receptor that binds simultaneously to the virus Spike protein and its main receptor ACE2, perhaps enhancing virus attachment. It also enhances the surface display of ACE2 in murine cells that overexpress human ACE2 (Carlos et al., 2021). A similar role of GRP78 was described for the replication cycle of Japanese Encephalitis virus (Nain et al., 2017) and Zika virus (Khongwichit et al., 2021). Our study does not rule out such a scenario, but GRP78 on the cell surface would not explain its synergistic role in Nirmatrelvir-treated cells. GRP78 is found both in the ER and on the cell surface (Ni et al., 2011), and the distribution between these compartments is subject to various regulatory mechanisms (Casas, 2017; Ni et al., 2011; Vig et al., 2019; Zhang et al., 2010). GRP78 contains a KDEL motif at its carboxyterminal end, typically mediating ER retention (Zhang et al., 2010). This argues in favor of an intracellular function of GRP78 during virus replication. This view is further supported by our observation that a GRP78 inhibitor still synergizes with Nirmatrelvir to block the infection cycle *after* having allowed virus entry.

If virus attachment does not represent the major contribution of GRP78 to SARS-CoV-2 replication in the context of Nirmatrelvir treatment, which other function of GRP78 is supporting the virus? GRP78 facilitates the import of proteins into the ER after their translation; it acts as a chaperone to assist proper folding of ER-localized proteins; it mediates retrograde transport of misfolded proteins from the ER to allow proteasomal degradation in the cytosol (ER-associated degradation, ERAD); and it triggers ER-stress signalling when dissociated from its protein clients IRE1, PERK, and ATF6 (Ibrahim et al., 2019; Ma and Hendershot, 2004; Zhu and Lee, 2015). Our assays cannot easily distinguish these possibilities. However, it is tempting to speculate that GRP78 can assist non-cleaved viral precursor polypeptides in relocating into the ER with their NSP4 and NSP6 subunits, and to assist in their folding within the ER. Along this line, an interaction between NSP6 and GRP78 was recently reported (Jiao et al., 2023). Proper location of the NSP precursor, in turn, would facilitate the autocleavage by the NSP5 subunit, even in the presence of non-saturating levels of Nirmatrelvir. With diminished GRP78 function, this relocation and folding of the precursor polypeptide may well be impaired. In such a situation, cleavage would be disturbed by improper folding as well as by Nirmatrelvir, explaining the synergistic inhibition of NSP5-driven autocleavage.

The role of ER-stress in SARS-CoV-2 replication is controversially debated. Thapsigargin, a drug that causes ER-stress, was reported to interfere with virus replication (Shaban et al., 2021); on the other hand, ER-stress can augment the synthesis of GRP78 by enhancing the transcription of its gene (Hetz et al., 2020; Lee, 2014; Shiu et al., 1977; Yoshida et al., 2000). However, it remains possible that, as part of interferon-induced GRP78 synthesis, e.g. in the context of infected tissue rather than cultured cells, infected cells may become less susceptible to Nirmatrelvir. This would further argue in favor of a clinical application of GRP78 inhibitors to improve antiviral treatment.

Inhibitors of viral proteases are also used to treat infections with Hepatitis C virus (HCV). It is conceivable that the efficacy of these inhibitors might also be enhanced by GRP78 inhibition. Indeed, the HCV precursor polypeptide, i.e. the substrate of protease cleavage, is also ER-associated (Pene et al., 2009; Reed and Rice, 2000). Thus, interfering with the ER-chaperone function might well synergize with protease inhibitors and leave behind uncleaved as well as unfolded precursor proteins in the context of HCV infections, too.

It would be desirable to apply drug synergies between antiviral protease inhibitors and pharmacological GRP78 antagonists in the clinics for improved treatment of infectious diseases. However, so far, GRP78 inhibitors have not been approved for clinical use. HA15 was at least tested in animals over several weeks, assessing its therapeutic impact on a malignant pleural mesothelioma model. Here, the animals survived such treatment without visible toxic effects (Xu et al., 2019). This raises hope that this or other GRP78 inhibitors can be developed for clinical use.

Nirmatrelvir, an M^Pro^ inhibitor, is currently used for treating COVID-19, but its clinical application is limited by two key factors. First, it requires co-administration with Ritonavir to inhibit liver metabolism, complicating treatment due to Ritonavir’s interactions with numerous other drugs, necessitating dose adjustments. Second, even with Ritonavir, Nirmatrelvir remains effective only in a subset of cases. The addition of a GRP78 inhibitor represents a promising strategy to overcome these limitations. By combining it with Nirmatrelvir, the dependency on Ritonavir could be reduced or even eliminated, as lower doses of Nirmatrelvir may suffice. Moreover, this combination may enhance overall therapeutic efficacy, offering a more effective and streamlined treatment option for COVID-19.

## MATERIALS AND METHODS

### Cell culture, drug treatment and transfection with siRNA

#### Cell culture

To maintain Vero E6 cells, Dulbecco’s modified Eagle’s medium (DMEM with GlutaMAX^TM^, Gibco) was used. DMEM was supplemented with 10% fetal bovine serum (Merck), L-Glutamine (200 mM), 50 units/mL penicillin, 50 µg/mL streptomycin (Gibco), 2 µg/mL tetracycline (Sigma) and 10 µg/mL ciprofloxacin (Bayer). Calu-3 cells were maintained in DMEM/F-12 GlutaMAX^TM^ (Thermo Fisher, 10565018), supplemented with 10% FCS (Merck), 1% Sodium pyruvate (PanBiotech, P04-43100), 1% non-essential amino acids solution (100x MEM NEAA, Thermo Fisher, 11140035), L-Glutamine (200 mM), 50 units/mL penicillin and 50 µg/mL streptomycin (Gibco). Both cell lines were maintained at 37°C in a humidified cell incubator with 5% CO_2_.

#### Drug treatment

20,000 Vero E6 cells/well were seeded into 24-well-plates using DMEM supplemented with 10% fetal bovine serum (FBS) and incubated for 24 h at 37°C and 5% CO_2_. Afterwards, the medium was changed to 2% FBS and the cells were infected with the Wuhan-like strain of SARS-CoV-2 (Stegmann et al. 2021) at MOI 0.02. 2 hpi, the cells were treated with Nirmatrelvir (Nirma, PF-07321332, MedChemExpress, HY-138687) and HA15 (MedChemExpress, HY-100437) for 48 h at the concentrations indicated in the figure legends. For time courses, 50,000 Vero E6 cells/well were seeded into 24-well-plates using DMEM supplemented with 10% fetal bovine serum (FBS) and incubated for 24 h at 37°C and 5% CO_2_. After 24 h, the medium was removed, and the cells were infected with the Wuhan-like strain of SARS-CoV-2 at MOI 2 in 2% FBS. 2 hpi, the inoculum was removed and the cells were washed three times with PBS. Afterwards, the cells were treated with Nirmatrelvir and HA15 at the concentrations indicated in the figure legends.

In the case of treating the cells with Celastrol (Sigma-Aldrich, C0869) or Main protease inhibitor 11a (Sigma-Aldrich, SML2877), 20,000 cells/well were incubated for 8 h at 37°C and 5% CO_2_. Then, the medium was changed to 2% FBS and the cells were treated with Celastrol or Main protease inhibitor 11a. After 24 h, the cells were infected with the Wuhan-like strain of SARS-CoV-2 and incubated for additional 48 h.

For the infection with the Omicron variant B.1.1.529 at MOI 0.1, 20,000 Vero E6 cells/well were seeded into 24-well-plates using DMEM supplemented with 10% fetal bovine serum (FBS) and incubated for 24 h at 37°C and 5% CO_2_. After 24 h, the 10% FBS medium was removed and the cells were infected with the Omicron variant B.1.1.529 in 2% FBS for 2 h.

Afterwards, the inoculum was removed, and the cells were washed three times with PBS and incubated with Nirmatrelvir and HA15 for 48 h.

When using Calu-3 cells, 80,000 cells/well were seeded into 24-well-plates using DMEM/F-12 GlutaMAX^TM^ supplemented with 10% fetal bovine serum (FBS) and incubated for 8 h at 37°C and 5% CO_2_. Afterwards, the medium was changed to 2% FBS and the cells were treated with Nirmatrelvir and HA15. After 24 h, the cells were infected with the Wuhan-like strain of SARS-CoV-2 and incubated for additional 48 h.

#### Transfection with siRNA

100,000 Vero E6 cells/well were reverse transfected with 22 nM siGRP78 (Thermo Fisher, s6979) or a pool of scrambled siRNAs (Thermo Fisher, 4390844; 4390847) using Lipofectamine 3000 (Thermo Fisher, L3000015). 24 h after the transfection, the cells were treated as described and harvested after 48 h.

| siRNA | Source |
| --- | --- |
| siGRP78 | Thermo Fisher Scientific: HSPA5 siRNA Human Coding Silencer® Select, s6979 |
| Scrambled siRNA #1 | Thermo Fisher Scientific, *Silencer*<sup>TM</sup> Select Negative Control No. 1 siRNA, 4390844 |
| Scrambled siRNA #2 | Thermo Fisher Scientific, *Silencer*<sup>TM</sup> Select Negative Control No. 2 siRNA, 4390847 |

### Virus titration

3,000 Vero E6 cells/well were first seeded into a 96-well plate using DMEM medium supplemented with 10% FBS and incubated for 8 h at 37°C and 5% CO_2_. After 24 h of culturing cells in medium containing 2% FBS, the cells were infected with SARS-CoV-2 in quadruplicates using ten-fold serial dilutions. At 48 hpi, the cytopathic effect was assessed by translucent microscopy. To determine the TCID_50_/mL, the plates were fixed with 4% PFA, permeabilized with 0.5% Triton X-100, stained with DAPI (Merck Sigma-Aldrich, D9542; 1:3000) and an antibody to Nucleoprotein (Sino Biological, 40143-R019; 1:8000) and imaged using a Celigo Imaging Cytometer. The titer was determined according to Spearman and Kärber (Kärber, 1931).

### Cytotoxicity assay

20,000 Vero E6 cells were seeded in 96-well plates and treated with Nirmatrelvir and/or HA15, for 72 h. The quantification of lactate dehydrogenase (LDH) release into the cell culture medium was performed through bioluminescence utilizing the LDH-Glo^TM^ Cytotoxicity Assay (Promega). To determine the maximum LDH release, 10% Triton X-100 was added to untreated cells for 15 min. The medium background (no-cell control) served as a negative control. Percent cytotoxicity reflects the proportion of LDH released into the media in comparison to the total LDH amount in the cells and was calculated using the following formula:

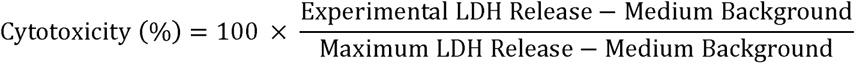

### RNA preparation and qRT-PCR

The cell culture supernatant containing SARS-CoV-2 was combined in a 1:1 ratio with the Lysis Binding Buffer from the MagNA Pure LC Total Nucleic Acid Isolation Kit (Roche) to inactivate the virus. Subsequently, viral RNA was extracted using Trizol LS, chloroform, and isopropanol. Following an ethanol wash of the RNA pellet, the purified RNA was resuspended in nuclease-free water. To assess the viral RNA yield, quantitative reverse transcription and polymerase chain reaction (qRT-PCR) was conducted following a previously established assay involving a TaqMan probe (Corman et al., 2020). The qRT-PCR utilized specific oligonucleotides targeting a genomic region corresponding to the envelope protein gene (26,141–26,253). The quantification of SARS-CoV-2 RNA during untreated infection was set as the reference (100%), and the other RNA quantities were normalized accordingly.

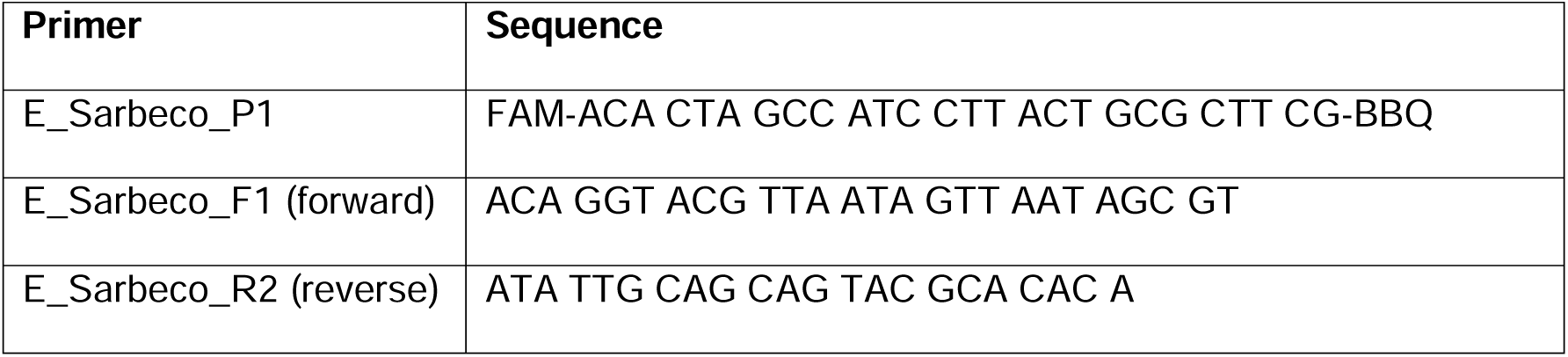

### Immunofluorescence

Vero E6 cells were seeded onto 8-well chamber slides (Nunc) and underwent treatment or infection as described in the corresponding figure legends. At 48 hpi, the cells were washed in PBS and fixed with 4% PFA in PBS for 1 h at room temperature. The cells were then permeabilized with 0.5% Triton X-100 in PBS for 30 min and blocked with 10% FBS in PBS for 20 min. The SARS-CoV-2 Spike protein (GeneTex, GTX 632604; 1:2000), Nucleoprotein (NP; Sino Biological, 40143-R019; 1:8000), NSP3 (Biozol, GTX135589; 1:1000) and NSP8 (Biozol, GTX632696; 1:1000) were stained overnight using primary antibodies.

The secondary antibodies, anti-rabbit IgG coupled to Alexa Fluor 546 (Thermo Fisher, A10040; 1:500), and anti-mouse IgG coupled to Alexa Fluor 488 (Thermo Fisher, A-21202; 1:500), as well as the nuclear stain 4′,6-diamidino-2-phenylindole (DAPI, Merck Sigma-Aldrich, D9542; 1:3000), were applied for 1 h at room temperature and then washed with PBS. The slides were mounted using Fluorescence Mounting Medium (DAKO, DakoCytomation, S302380-2). Fluorescence signals were detected using epifluorescence and confocal microscopy (Zeiss).

### Immunoblot analysis

After washing the cells once with PBS, they were harvested in radioimmunoprecipitation assay (RIPA) lysis buffer (20 mM TRIS-HCl (pH 7.5), 150 mM NaCl, 10 mM EDTA, 1% Triton X-100, 1% deoxycholate salt, 0.1% SDS, 2 M urea) with freshly added protease inhibitors (Merck Sigma-Aldrich, 5056489001). Following sonication, protein extracts were quantified using the Pierce BCA Protein assay kit (Thermo Fisher, 23227). After equalizing protein concentrations, the samples were incubated at 95°C in Laemmli buffer for 5 min and then separated through SDS-PAGE.

The separated proteins were transferred onto nitrocellulose or PVDF membranes, followed by blocking in 5% (w/v) non-fat milk in TBS containing 0.1% Tween 20 for 1 h. The membranes were incubated with primary antibodies overnight at 4°C, followed by incubation with peroxidase-conjugated secondary antibodies (Jackson Immunoresearch, 715-036-150, 715-036-152) for 1 h at room temperature. Detection of the proteins was performed using either Immobilon Western Substrate (Millipore, WBKLS0500) or Super Signal West Femto Maximum Sensitivity Substrate (Thermo Fisher, 34095). The following primary antibodies were used.

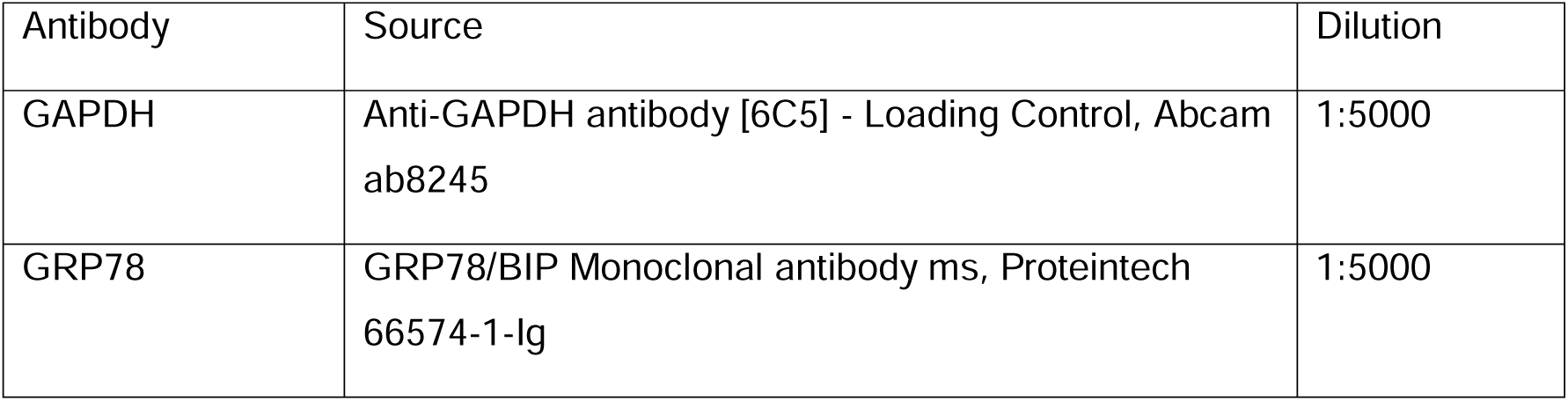
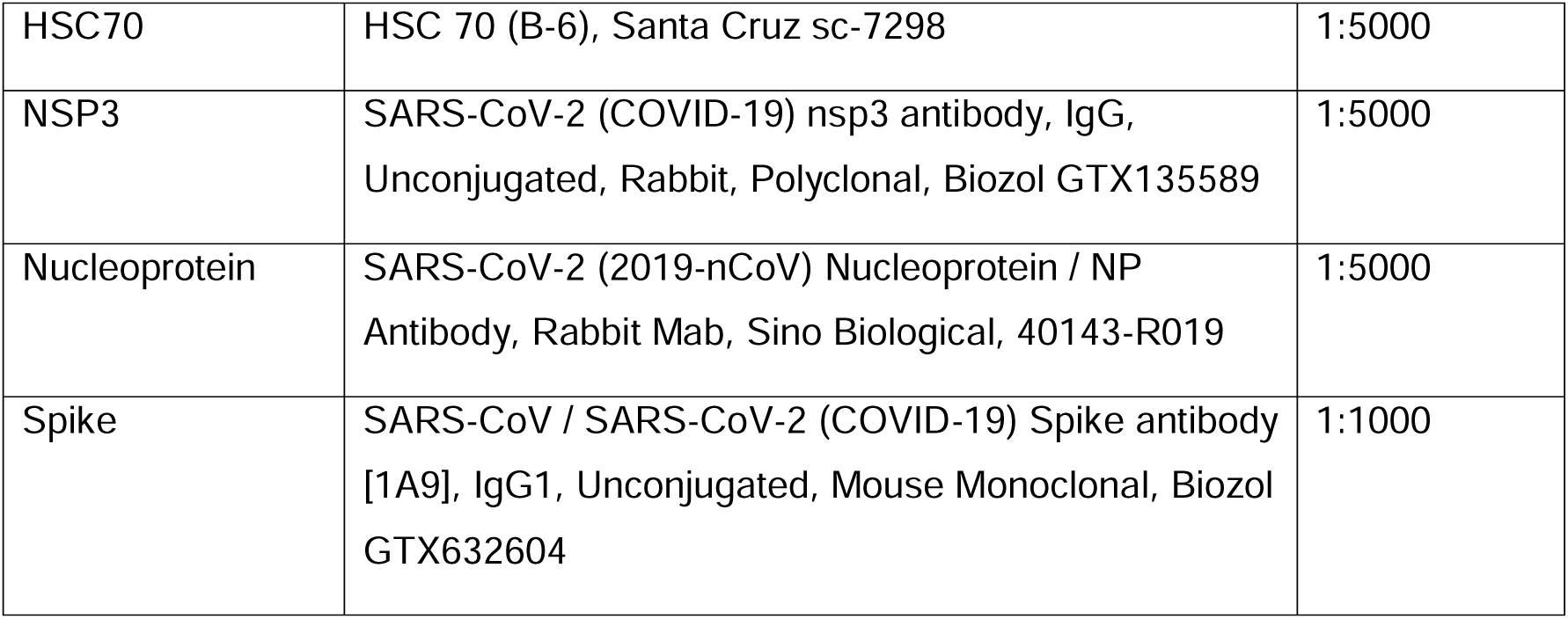

### Statistical analyses

Statistical analyses were performed using GraphPad Prism 10. Due to the exponential growth kinetics observed in the experiments, a one-way ANOVA was conducted on the geometric mean, followed by Dunnett’s multiple comparisons test.

### Animal experiments

#### Ethics statement

Hamster experiments were carried out according to the German Regulations for Animal Welfare after obtaining the necessary approval from the authorized ethics committee of the State Office of Agriculture, Food Safety and Fishery in Mecklenburg – Western Pomerania (LALLF MV) under permission number 7221.3-1-049/20 and approval of the commissioner for animal welfare at the Friedrich-Loeffler-Institute (FLI), representing the Institutional Animal Care and Use Committee (IACUC).

#### Infection and treatment

Male Golden Syrian Hamsters (Mesocricetus auratus; RjHan:AURA; 80–100 g) were obtained from Janvier Labs (Saint Berthevin, France) and were housed in standard rodent IVCs Type III in groups of 3 to 4 animals under standardized conditions (22°C; 12/12h light cycle). As diet, rodent pellets and water were offered ad libitum. For acclimatization, rodents were housed under these conditions for one week prior to inoculation. The experiments were conducted in a BSL-3 animal facility. Animals were infected orotracheally with 1*10^4^ TCID_50_ SARS-CoV-2 Germany/BavPat1/2020 (BavPat1) (Wolfel et al., 2020) (GISAID accession EPI_ISL_406862) in a volume of 100 µL. The mock-infected animals received the same volume of cell culture medium.

Drugs were administered daily from one day prior to the infection until 6 days post infection (dpi). Nirmatrelvir was dissolved in Aqua bidest to a concentration of 125 mg/ml. Each animal received 250µl twice / day orally. Ritonavir was dissolved in Aqua bidest to a concentration of 50 mg/ml. Each animal received 250µl / day orally. HA15 was purchased as a 50 mg/ml solution, and each animal was administered 5 mg/day intraperitoneally.

Animals were clinically observed for 7 days with a daily sampling for virological analysis. After 7 days, animals were euthanized by inhalation of an isoflurane overdose followed by intracardial exsanguination and decapitation. Virological analysis of the collected samples was performed as described in (Blaurock et al., 2022).

#### Pathology

For histopathology, the left lung lobe was carefully removed, immersion-fixed in 10% neutral buffered formalin, embedded in paraffin, and 2-3-μm sections were stained with hematoxylin and eosin (HE). All slides were scanned with a Hamamatsu S60 scanner and analysed by a trained pathologist (TB) and reviewed by a certified pathologist (AB) in a blinded manner using NDPview.2 plus software (Hamamatsu Photonics, K.K., Japan). The extent of consolidation resulting from pneumonia was evaluated by quantifying the percentage of affected lung fields, which was assessed using a 500 × 500 μm grid. In addition, a semi-quantitative examination of lung tissue was performed to determine the presence of SARS-CoV-2 specific lesions observed in hamsters as given in Suppl. Table 1. These lesions included intra-alveolar, interstitial, peribronchial and perivascular inflammatory infiltrates, alveolar edema, necrosis of the bronchial epithelium, diffuse alveolar damage, vasculitis, activation of endothelium with immune cell rolling, as well as bronchial epithelial and pneumocyte type 2 hyperplasia. As given in Fig. S13B the sum of the values for alveolar, interstitial, peribronchial, perivascular infiltrates and vasculitis constituted an inflammation score, while the sum of the values for bronchial epithelial and pneumocyte type 2 hyperplasia formed a regeneration score and the values for perivascular infiltrates, vasculitis and immune cell rolling were summarized in a vascular score. Statistical analysis was performed using GraphPad Prism 9 software. The arithmetic means of pulmonary atelectasis and the medians of the inflammation, regeneration and vascular scores were used.

### Materials

#### Antibodies

| Antibody | Source | Identifier |
| --- | --- | --- |
| Alexa Fluor 488 donkey anti-mouse IgG | Thermo Fisher Scientific | A-21202 |
| Alexa Fluor 488 donkey anti-rabbit IgG (H+L) | Thermo Fisher Scientific | A32790 |
| Alexa Fluor 546 donkey anti-rabbit IgG | Thermo Fisher Scientific | A10040 |
| DAPI | Sigma-Aldrich | Cat# D9542 |
| GAPDH | Abcam | ab8245 |
| GRP78 | Proteintech | 66574-1-Ig |
| HSC70 | Santa Cruz | sc-7298 |
| SARS-CoV-2 NSP3 | Biozol | GTX135589 |
| SARS-CoV-2 NSP8 | Biozol | GTX632696 |
| SARS-CoV-2 Nucleoprotein | Sino Biological | 40143-R019 |
| SARS-CoV-2 Spike | Biozol | GTX632604 |

#### Chemicals, peptides, reagents

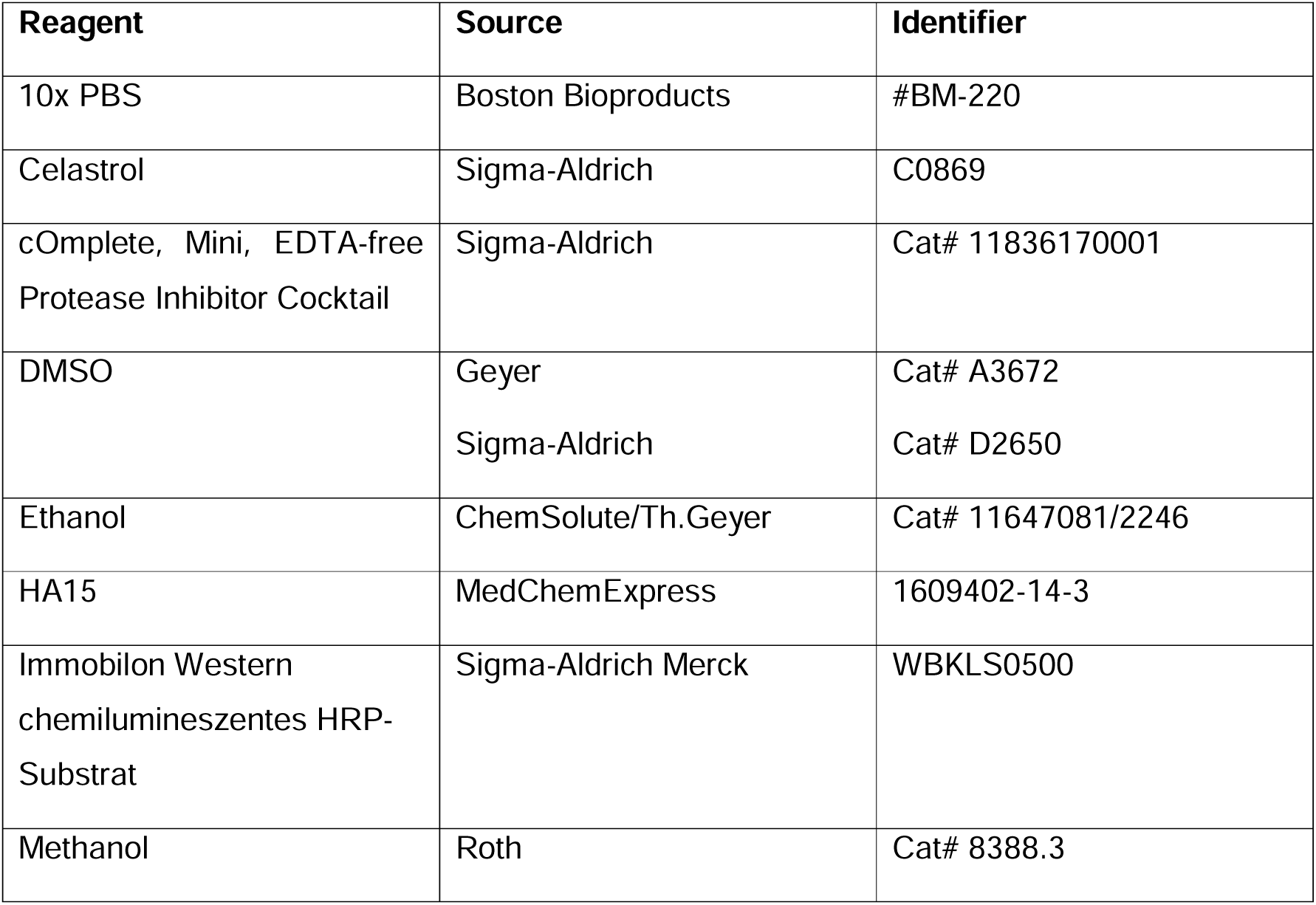
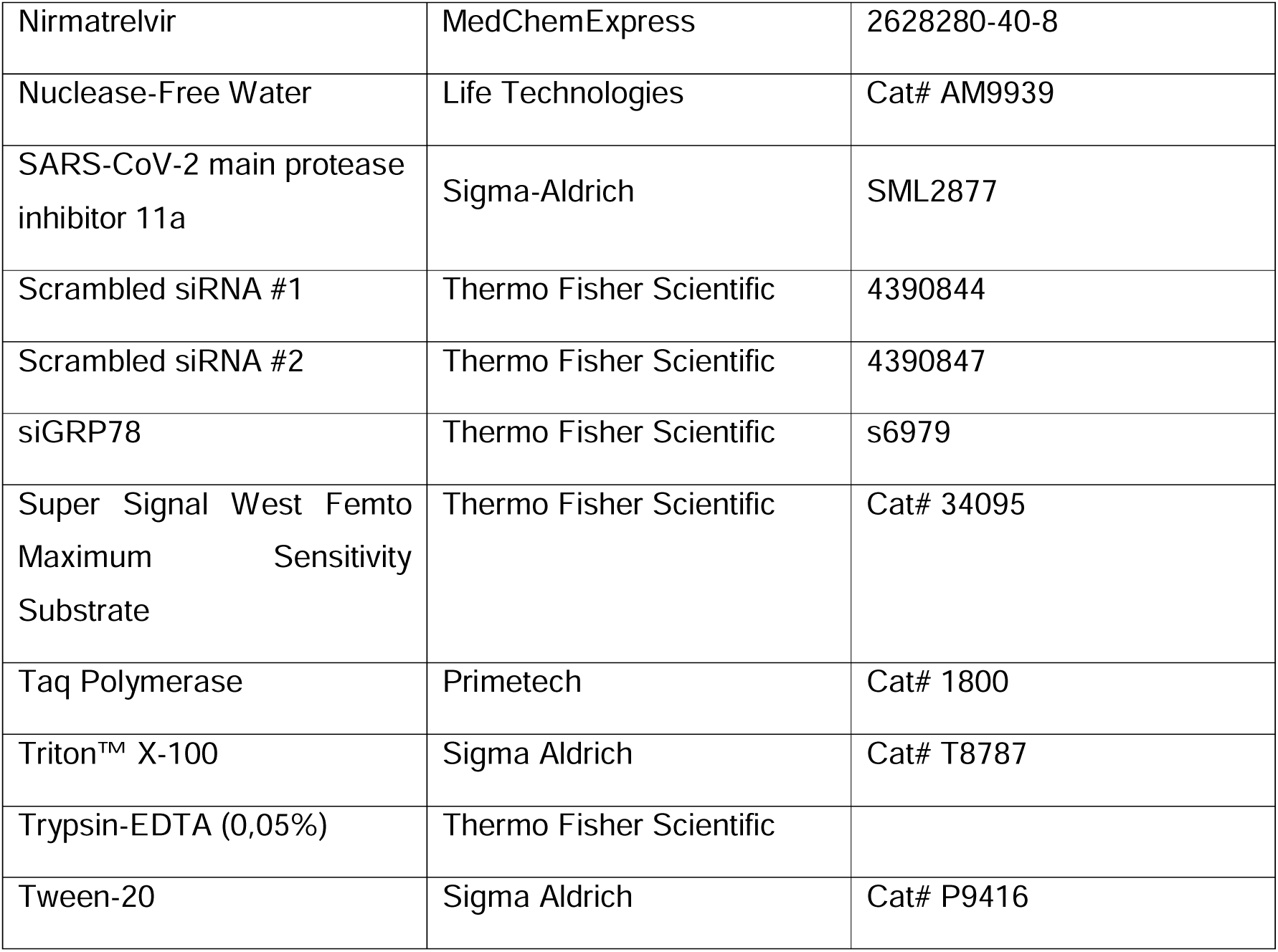

#### Commercial assays

| **Assay** | **Source** | **Identifier** |
| --- | --- | --- |
| LDH-Glo™ Cytotoxicity Assay kit | Promega | Cat# J2380 |
| Pierce BCA Protein Assay Kit | Thermo Fisher Scientific | Cat# 23227 |

#### Experimental models: Cell lines

| **Cell line** | **Source** | **Identifier** |
| --- | --- | --- |
| Calu-3 | ATCC | HTB-55 |
| Vero E6 | ATCC | CRL-1586 |

#### Experimental models: Animals

| Animal | Source | Identifier |
| --- | --- | --- |
| Male Golden Syrian Hamsters (Mesocricetus auratus; RjHan:AURA) | Janvier Labs (Saint Berthevin, France) | N/A |

#### Cell culture media

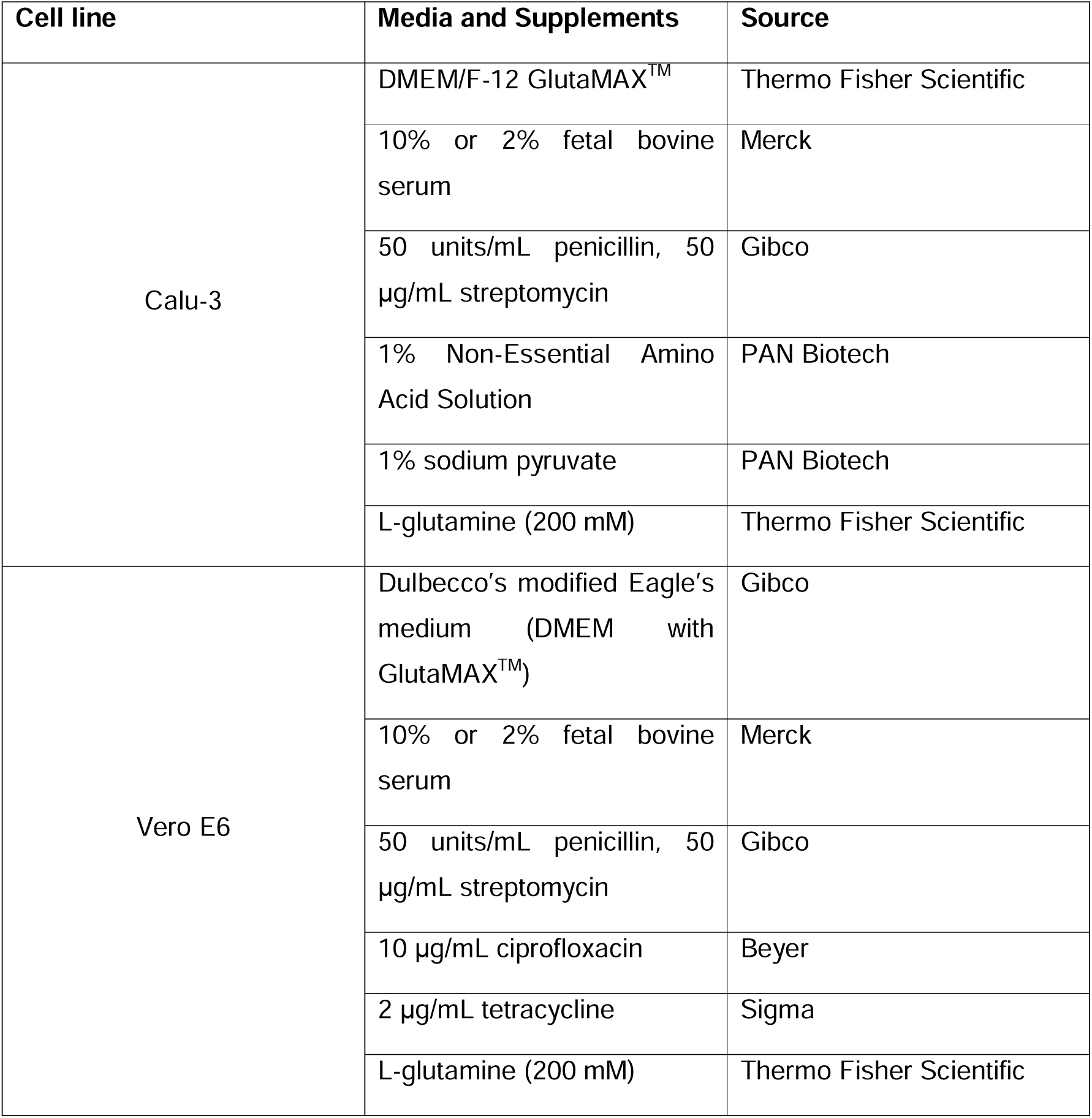

#### Oligonucleotides (metabion)

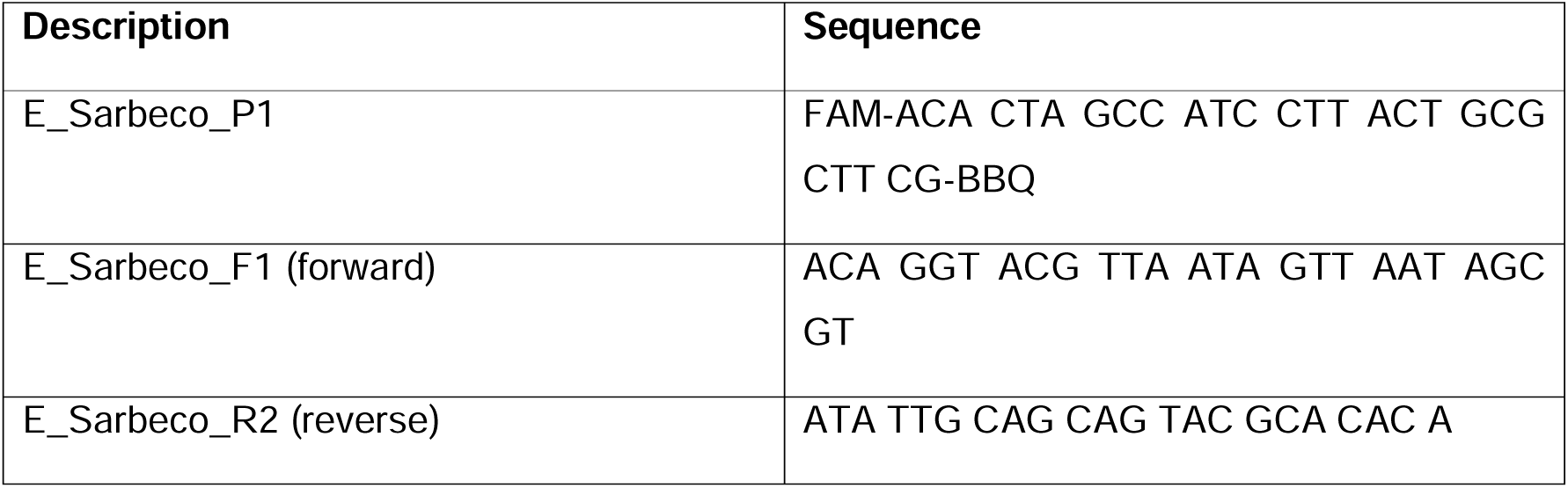

#### Software and algorithms

| Software and algorithms | Source | Identifier |
| --- | --- | --- |
| Image Lab v. 5.2.1 | Bio Rad | N/A |
| NDPview.2 plus software | Hamamatsu Photonics,<br>K.K., Japan | N/A |
| Prism v. 10.2.0 | GraphPad | N/A |
| SynergyFinder+ | SynergyFinder | https://synergyfinder.org/ |
| TCID50 calculator (v2.1 -<br>20-01-2017_MB) | *by Marco Binder; adapted<br>@ TWC | N/A |
| ZEN | Zeiss | N/A |

#### Virus Strains

| Description | Source | Identifier |
| --- | --- | --- |
| SARS-CoV-2 'wildtype',<br>Göttingen/Germany | Isolated from patient<br>(Stegmann et al., 2021) | N/A |
| SARS-CoV-2, omicron<br>variant, B.1.1.529 | Robert Koch Institute<br>(Berlin, Germany) | Ref-SKU: 009V-04437 |
| SARS-CoV-2<br>Germany/BavPat1/2020<br>(BavPat1) | Friedrich Loeffler Institute<br>(Germany) | GISAID: EPI_ISL_406862 |

## ACKNOWLEDGEMENTS

We gratefully acknowledge the technical assistance by Rebecca Soliwoda and Robin Brandt.

This work was funded by Volkswagenstiftung VW-9B785 and the Corona Forschungsnetzwerk Niedersachsen (CoFoNi). D.A.K. was supported by the Studienstiftung des deutschen Volkes and the Promotionskolleg funded by the Else Kröner-Fresenius foundation during this work. K.M.S. was supported by the Göttingen Graduate Center for Neurosciences, Biophysics, and Molecular Biosciences, GGNB.

---

**Figure S1.**
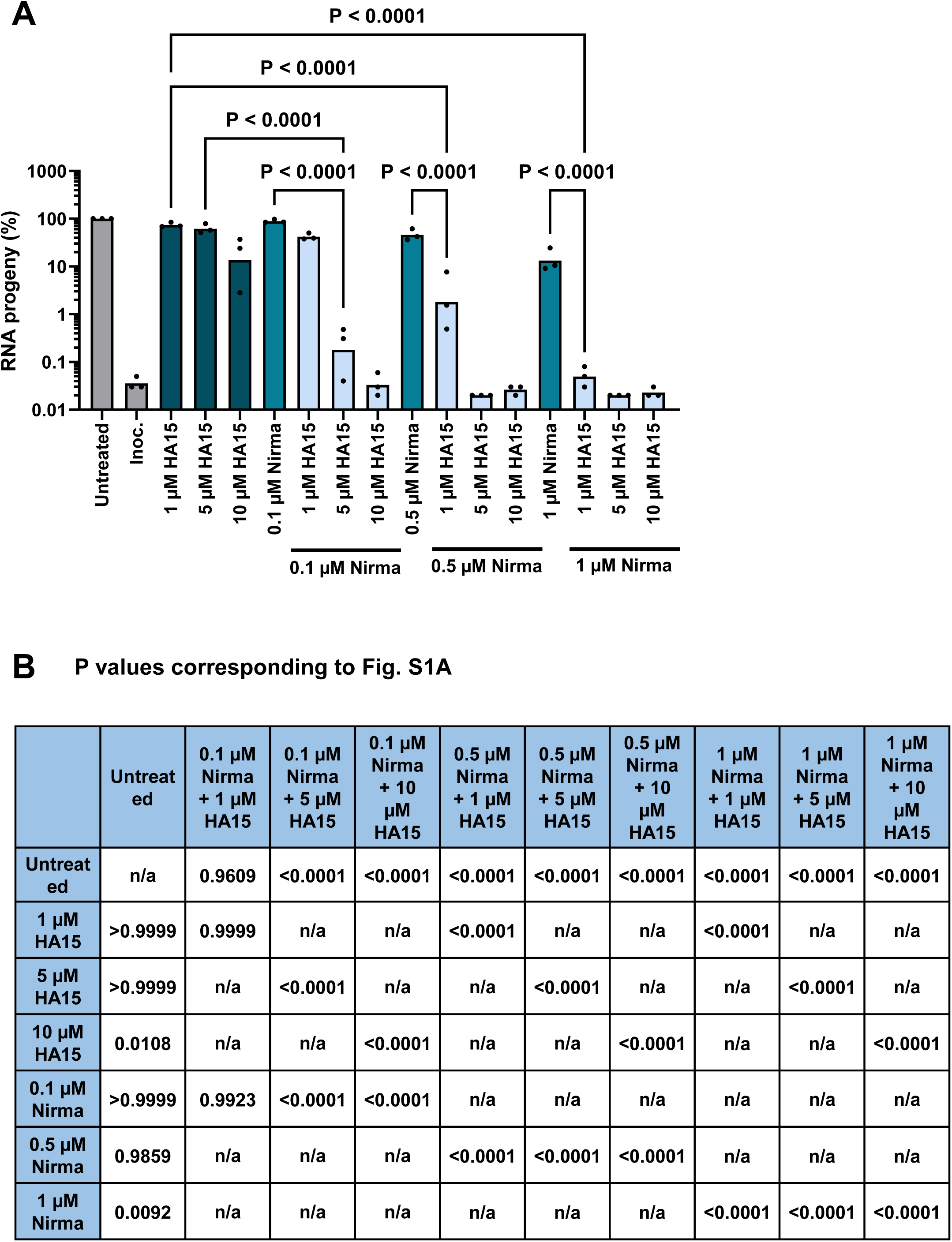
related to Fig. 1: Virus RNA quantification upon treatment with Nirmatrelvir and/or HA15. A. Viral RNA progeny corresponding to Fig. 1B. B. P values corresponding to Fig. S1A.

**Figure S2.**
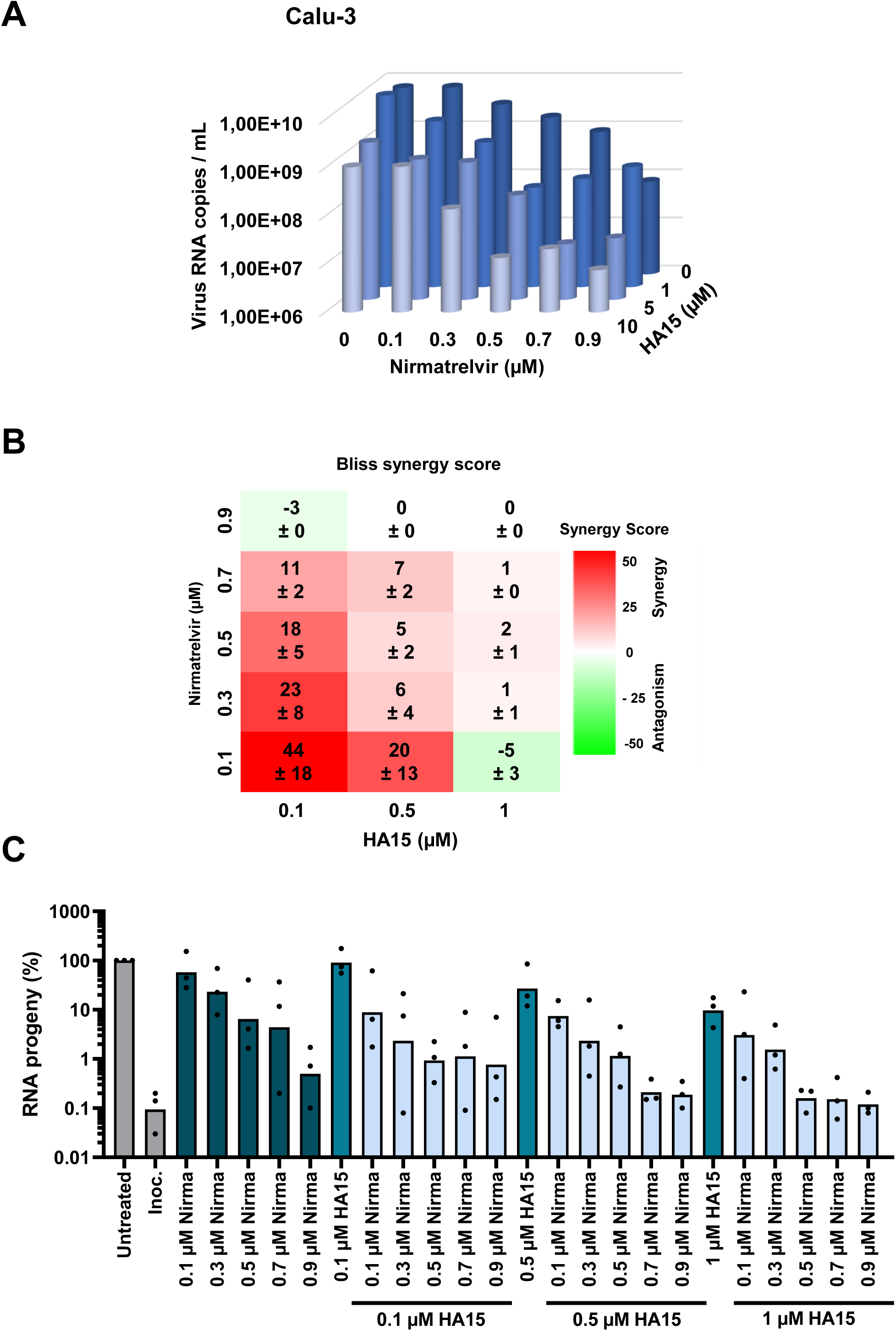
related to Fig. 1: SARS-CoV-2 replication in Calu-3 cells. A. Strongly reduced yield of virus RNA in Calu-3 cells. Calu-3 cells were treated and infected as in Fig. 1A. At 48 hpi, virus RNA in the supernatant was quantified (n = 3). B. The Bliss independence model was used to calculate the synergy scores. Bliss values > 10 indicate strong drug synergy (mean ± SEM). C. Viral RNA progeny corresponding to Fig. S2A.

**Figure S3.**
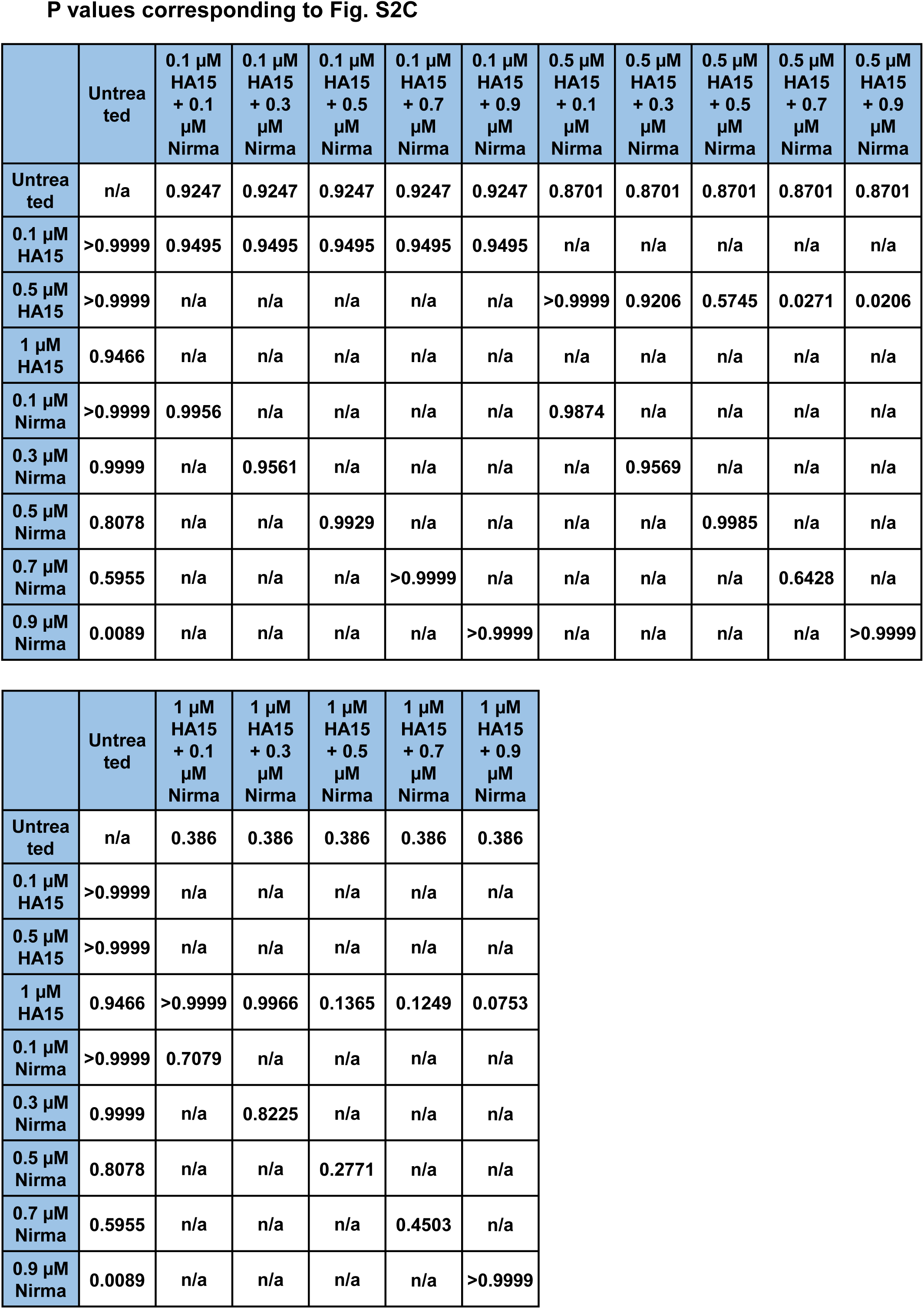
related to Fig. 1: Drug response in Calu-3 cells. P values corresponding to Fig. S2C.

**Figure S4.**
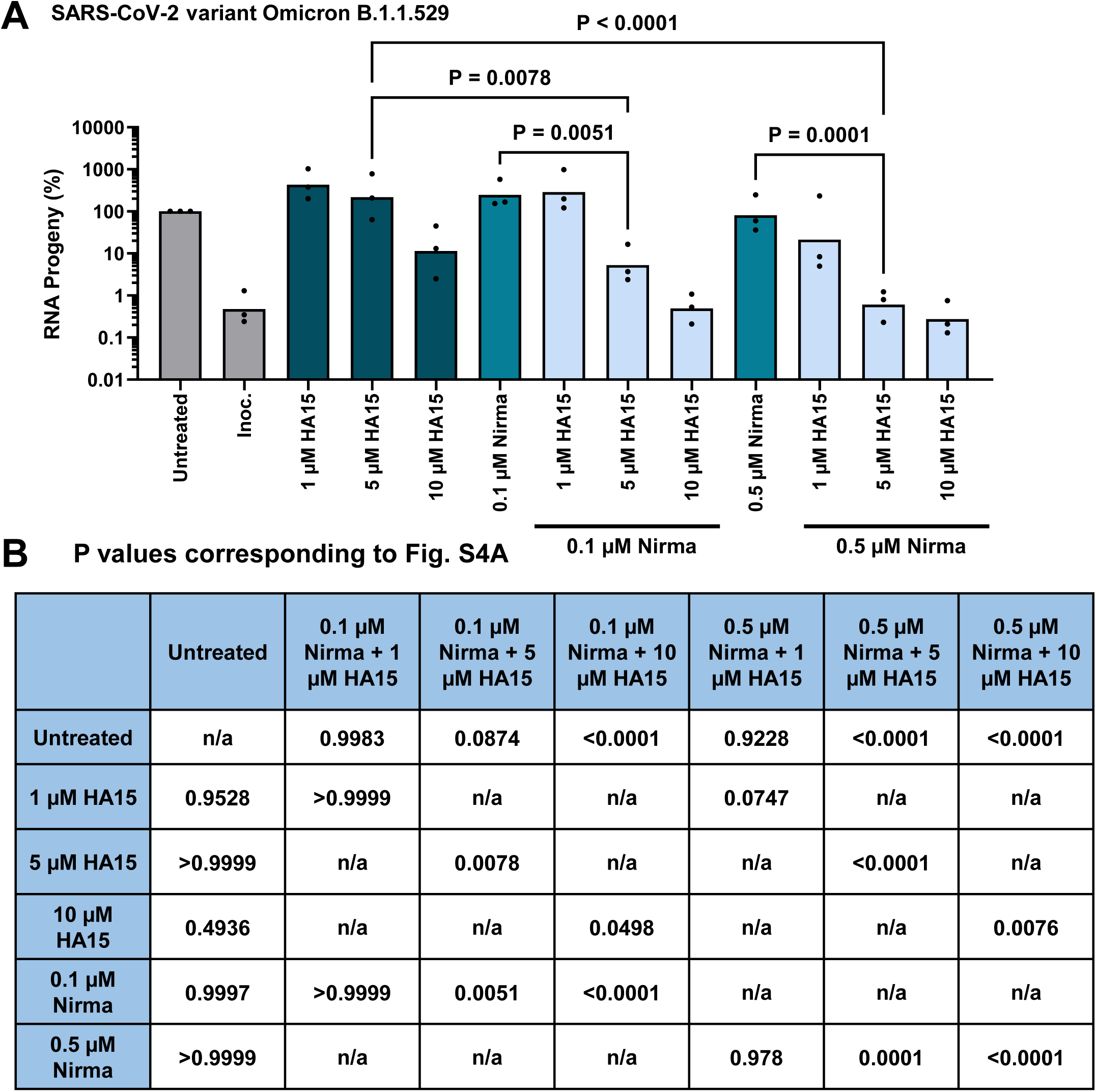
related to Fig. 3: SARS-CoV-2 Omicron, drug response. A. Viral RNA progeny corresponding to Fig. 3A. B. P values corresponding to Fig. S4A.

**Figure S5.**
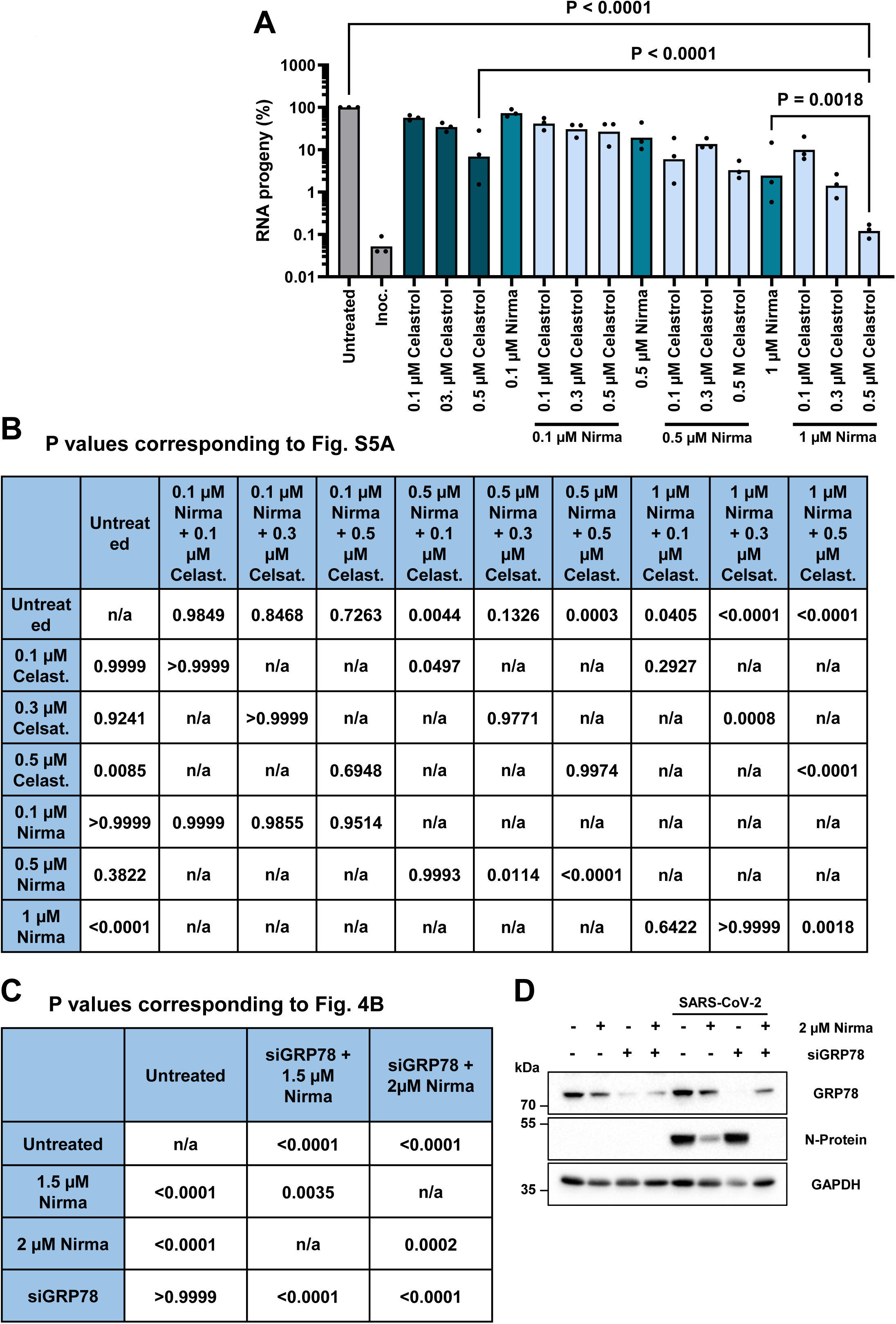
related to Fig. 4: Celasterol and GRP78 siRNA. A. Viral RNA progeny corresponding to Fig. 4A. B. P values corresponding to Fig. S5A. C. P values corresponding to Fig. 4B. D. Vero E6 cells were transfected with siRNAs to deplete GRP78 and treated as in Fig. 4C using a higher Nirmatrelvir concentration.

**Figure S6.**
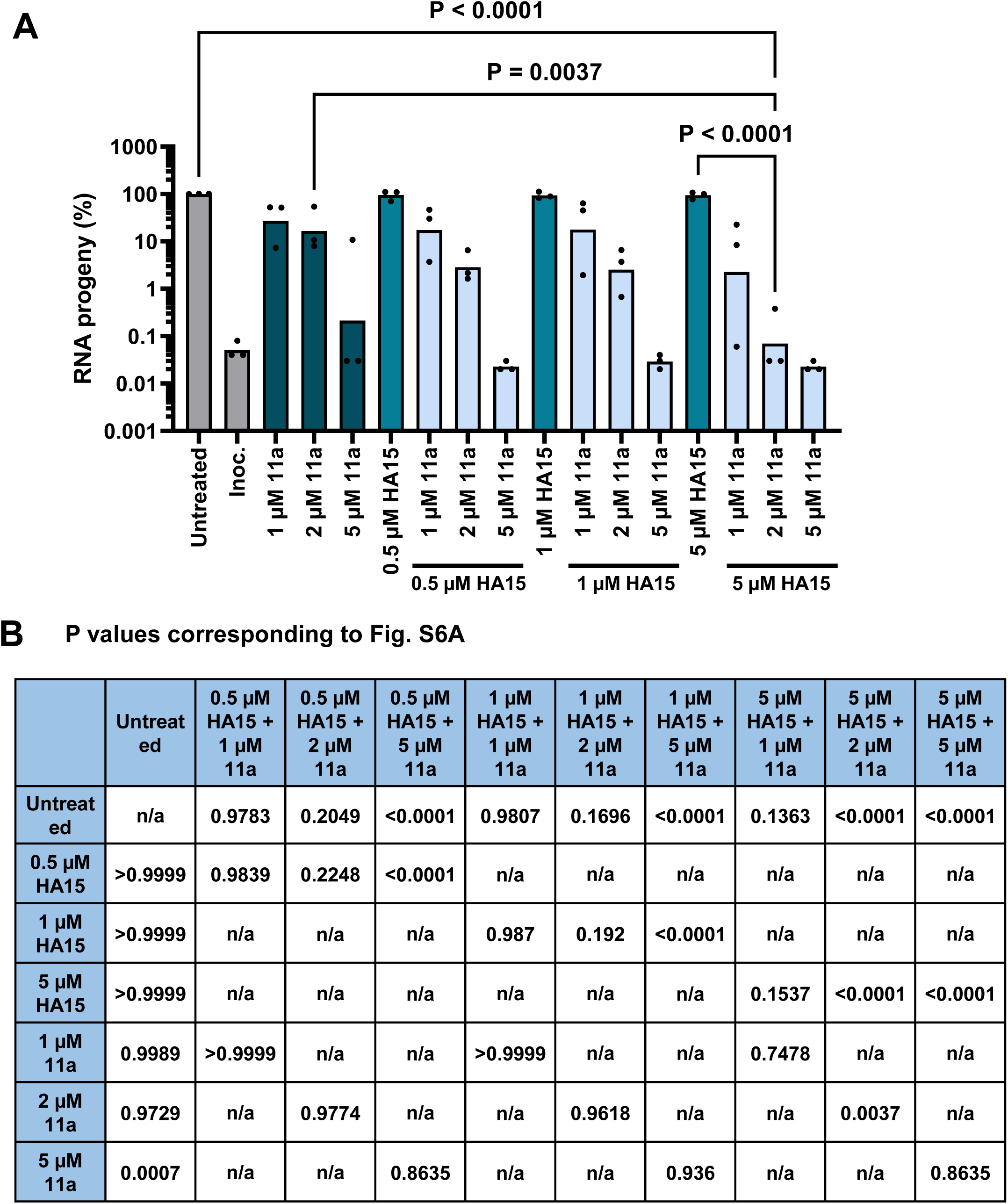
related to Fig. 4: M^Pro^ inhibitor 11a. A. Viral RNA progeny corresponding to Fig. 4D. B. P values corresponding to Fig. S6A.

**Figure S7.**
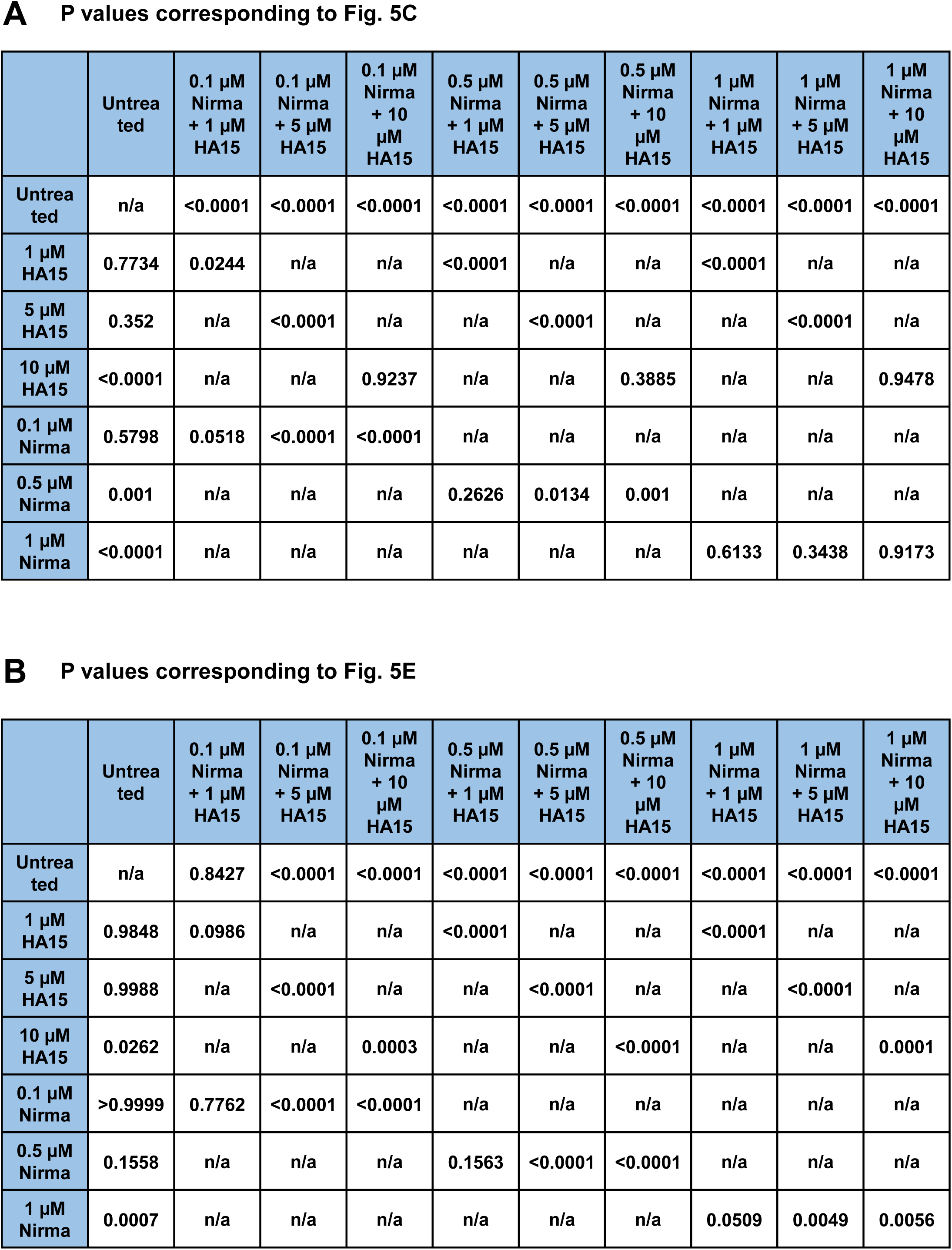
related to Fig. 5: Early accumulation of virus RNA. A. P values corresponding to Fig. 5C. B. P values corresponding to Fig. 5E.

**Figure S8.**
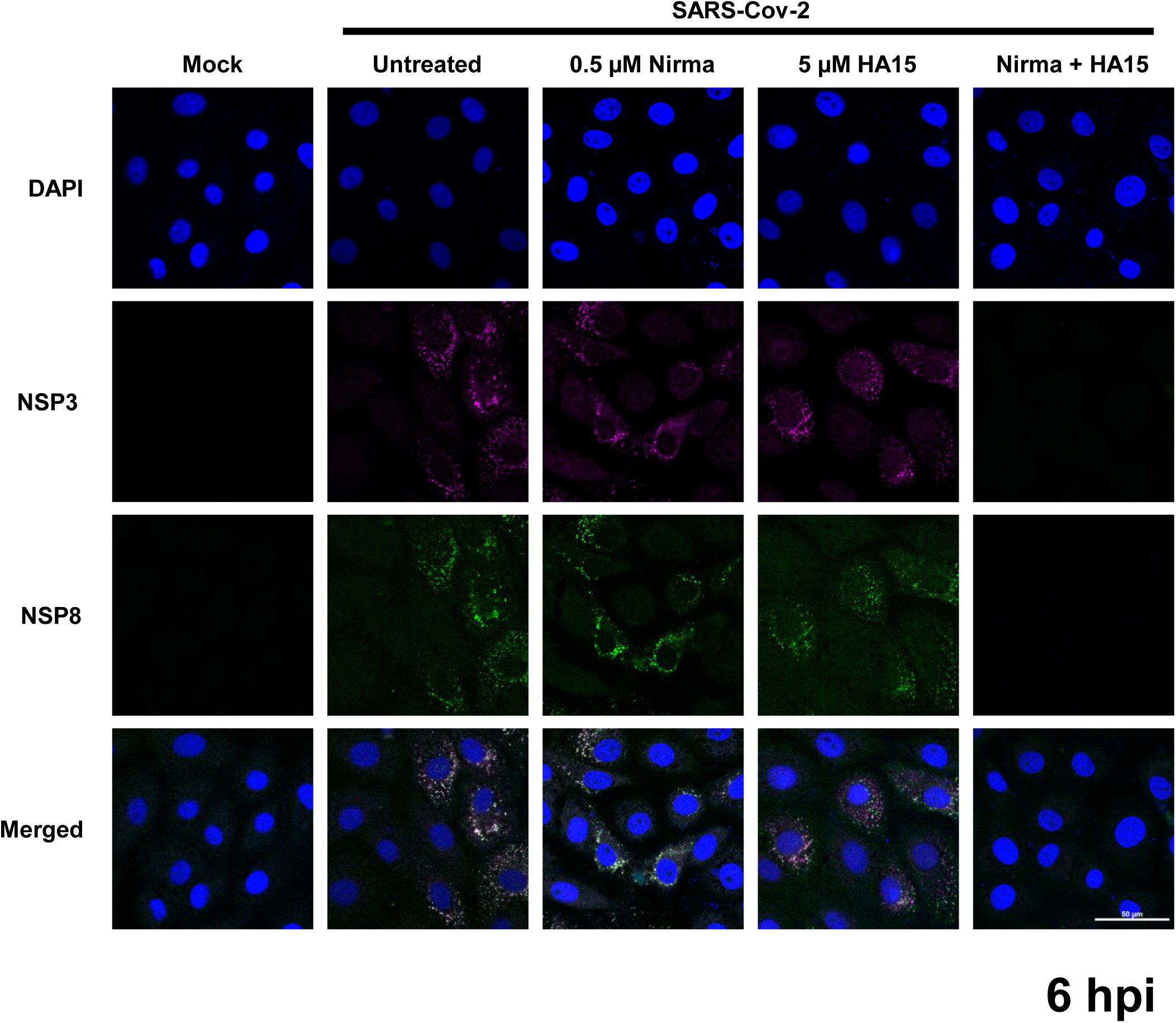
related to Fig. 6: Diminished accumulation of NSPs in the presence of Nirmatrelvir and HA15. Vero E6 cells were treated and 24 h later infected with SARS-CoV-2 at MOI 2. To analyse the cells using immunofluorescence microscopy, they were fixed and stained with antibodies to detect NSP3 and NSP8 during the early stages of the viral replication cycle, here 6 hpi. Scale bar, 50 µM.

**Figure S9.**
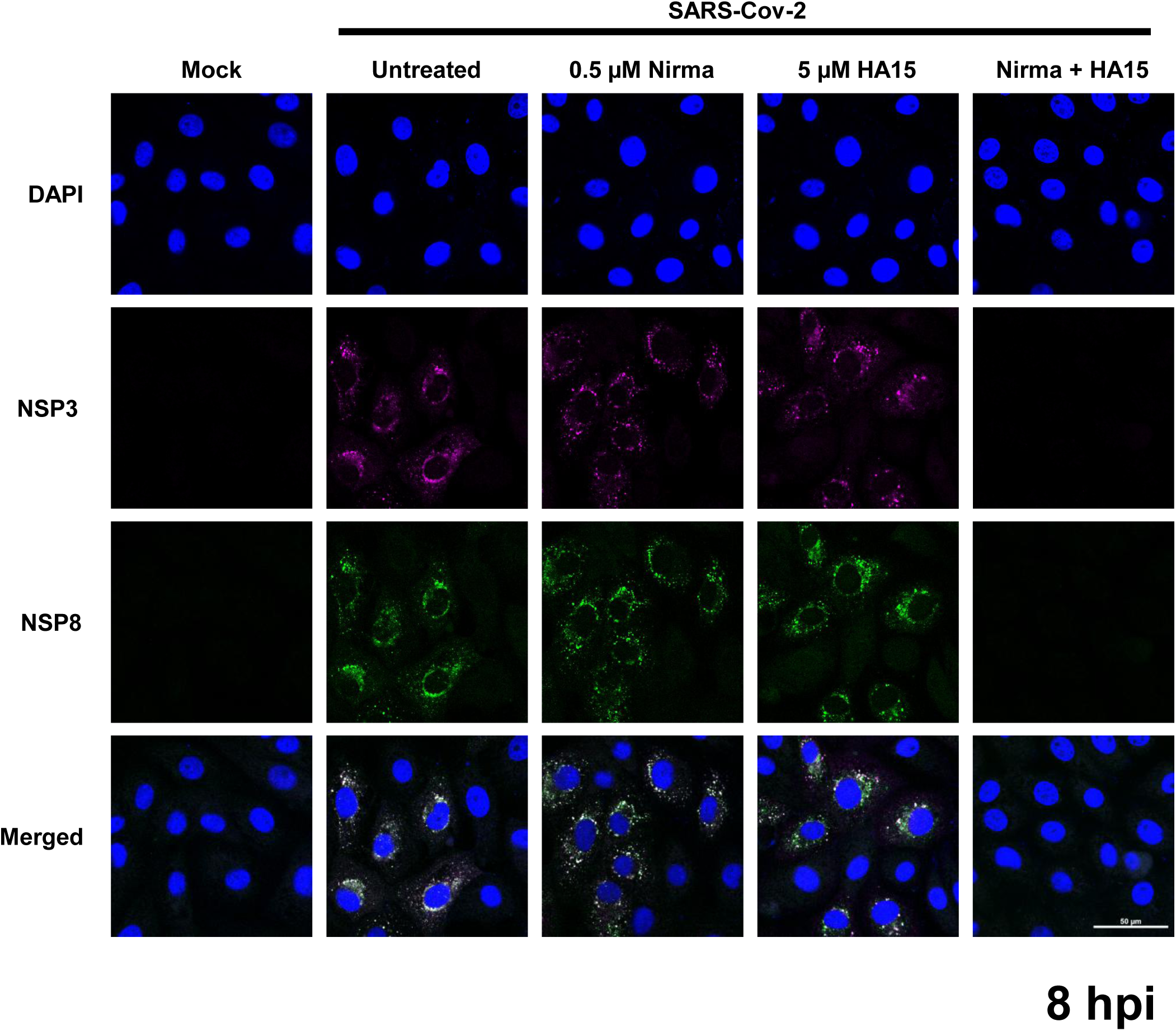
related to Fig. 6: NSP accumulation at 8 hpi. Vero E6 cells were treated and 24 h later infected with SARS-CoV-2 at MOI 2. To analyse the cells using immunofluorescence microscopy, they were fixed and stained with antibodies to detect NSP3 and NSP8 during the early stages of the viral replication cycle, here 8 hpi. Scale bar, 50 µM.

**Figure S10.**
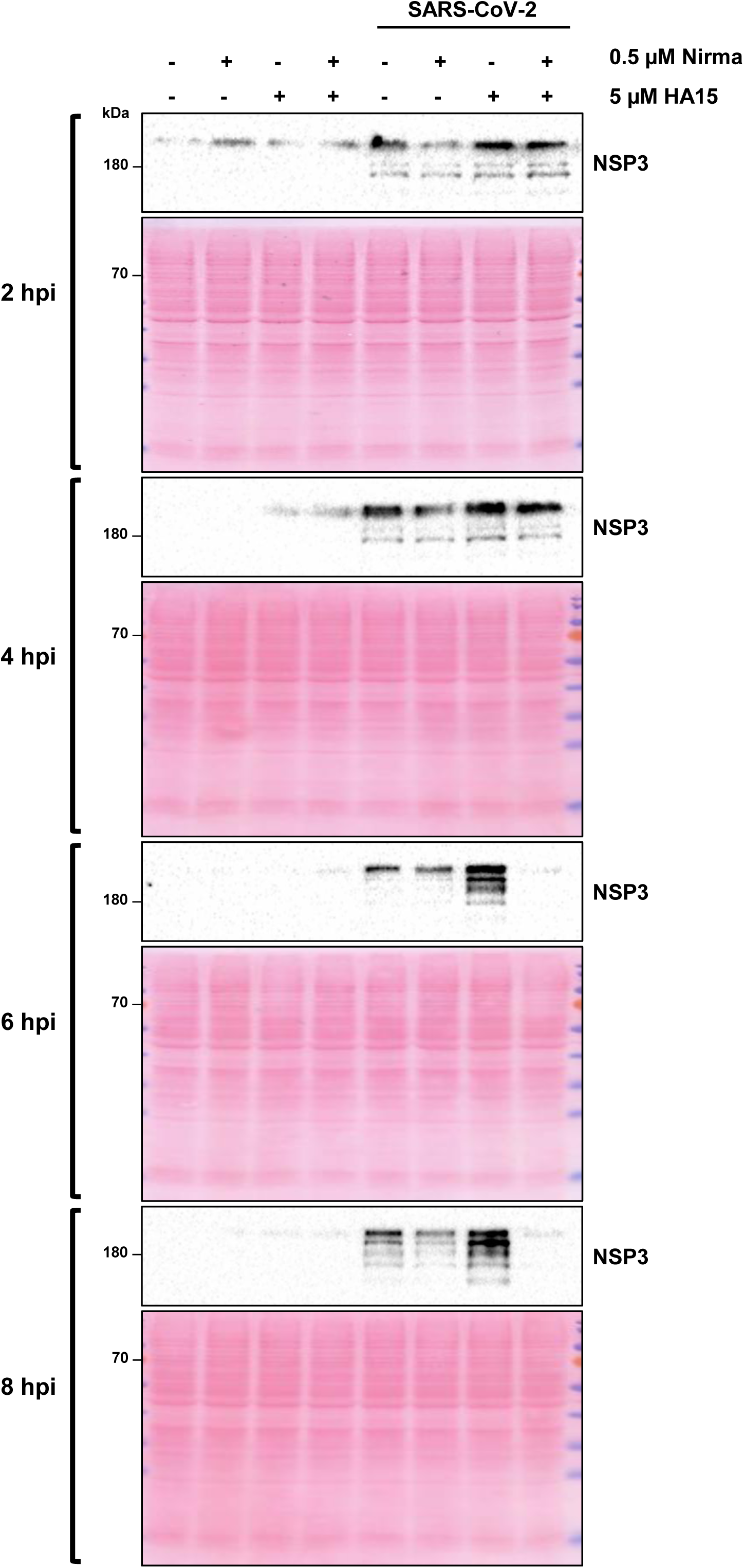
related to Fig. 6: Diminished accumulation of NSP3 in the presence of Nirmatrelvir and HA15. Vero E6 cells were treated and 24 h later infected with SARS-CoV-2 at MOI 2. The cell lysates were analysed by immunoblot. Nirmatrelvir and HA15 supressed almost completely the synthesis of NSP3, as early as 6 hpi.

**Figure S11.**
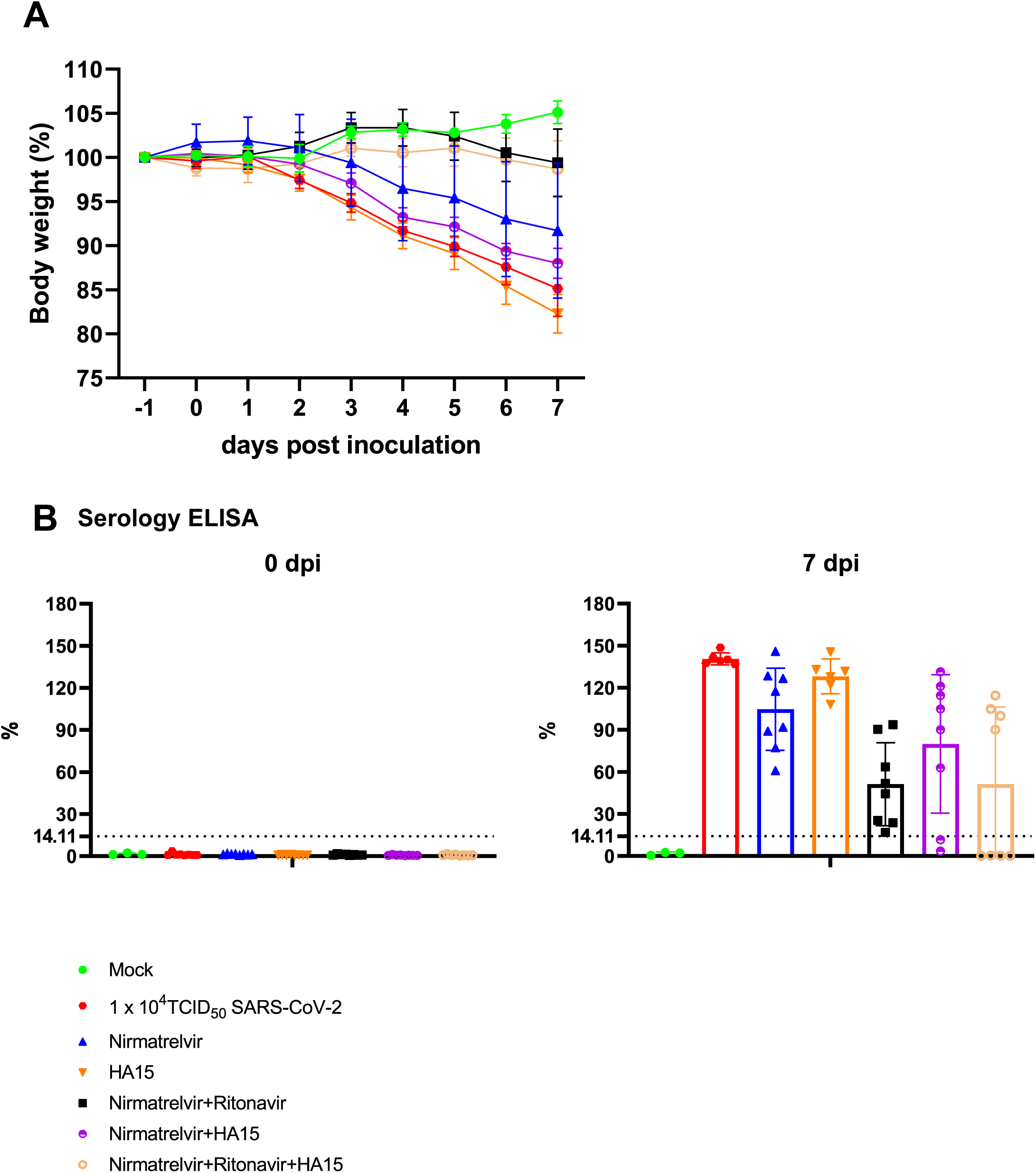
related to Fig. 7: Animal model, body weight and serum analysis. A. Upon treatment and infection as in Fig. 7A, the body weight of the animals were determined. B. Antibodies against the Spike protein of SARS-CoV-2 in the blood of the animals were quantified by Enzyme-Linked ImmunoSorbent Assay (ELISA) before infection and on the day of termination.

**Figure S12.**
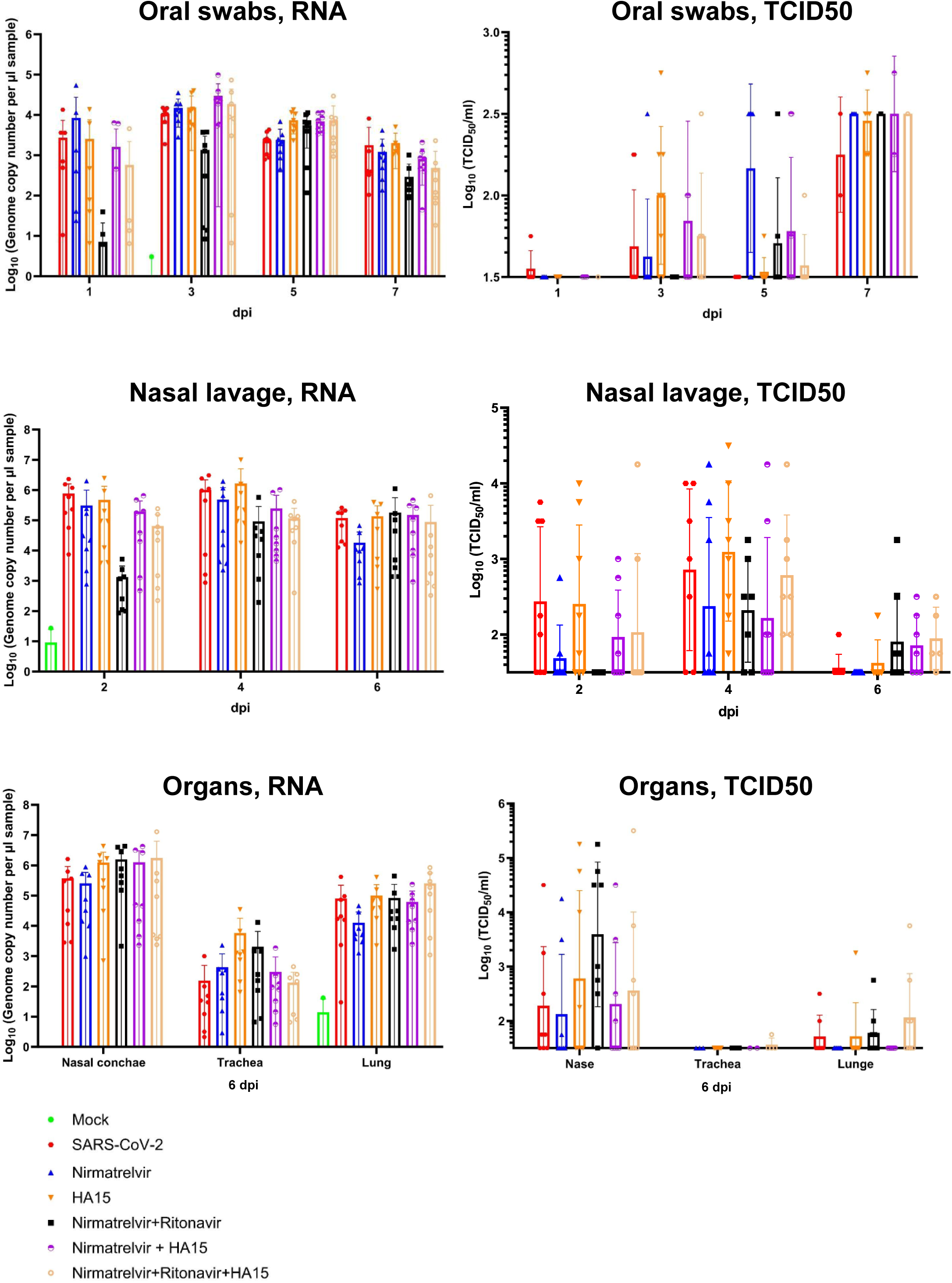
related to Fig. 7: Virus titers in the animals. Virus RNA levels (qRT-PCR) and virus titers (TCID_50_) in oral swabs and nasal lavage at the indicated time points (days post infection, dpi), and in the organs at the day of termination, determined as described previously (Schrell et al., 2025).

**Figure S13.**
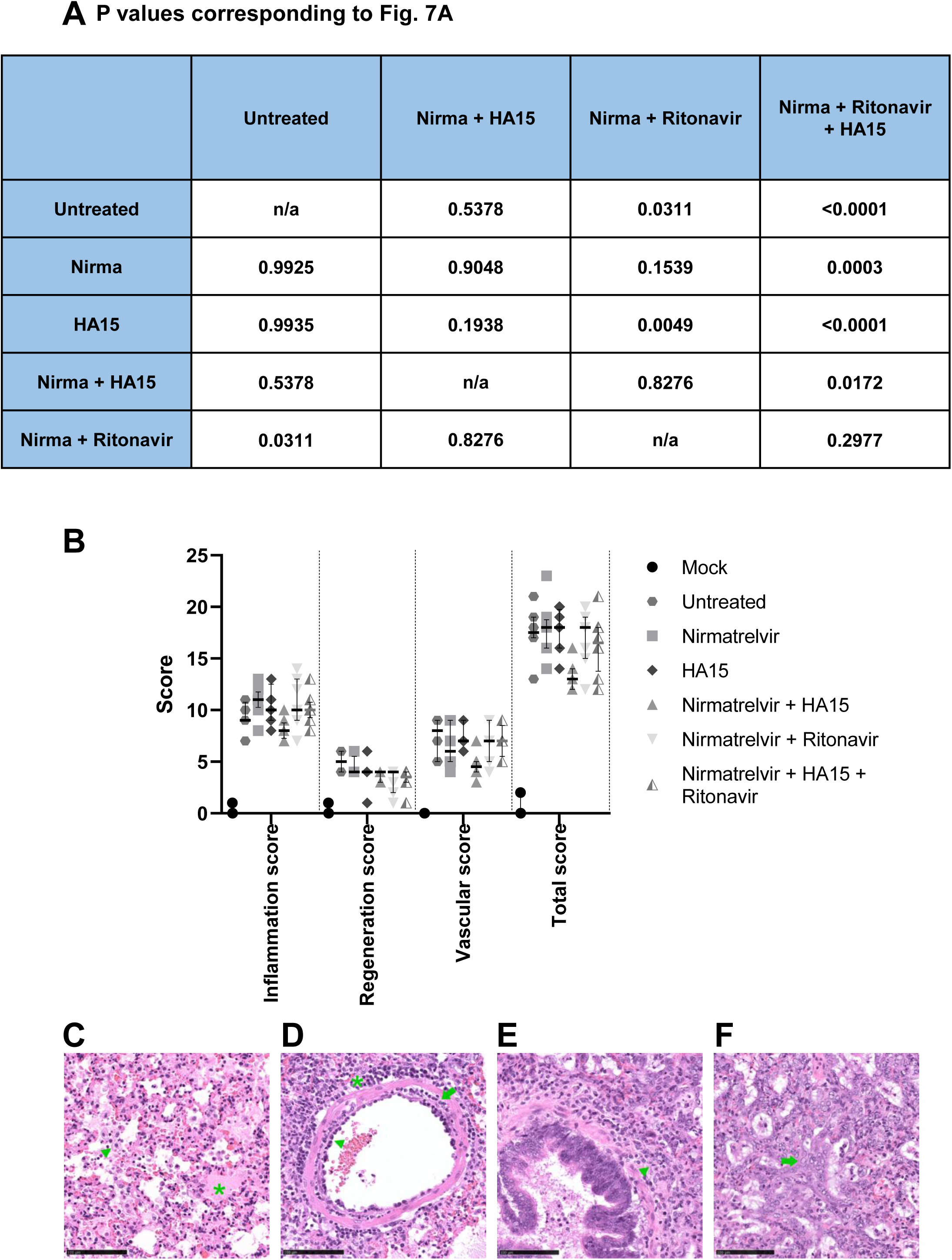
related to Fig. 7: Histopathology. A. P values corresponding to Fig. 7A. B. Histopathological scores. Median with interquartile range is shown. Kruskal–Wallis test with Dunn’s correction was performed. No statistical significance was found. C-F: SARS-CoV-2-related lesions 6 days post infection. C. Alveolar immune cell infiltration, mainly macrophages, (green arrowhead) and alveolar edema (green asterisk), Nirmatrelvir-HA15-Ritonavir treatment, found in 8/8 hamsters. Scale bar, 100 µm. D. Blood vessel with vasculitis (green arrow), perivascular immune cell infiltration, mainly macrophages and lymphocytes, (green asterisk) and rolling of immune cells (green arrowhead), Nirmatrelvir-Ritonavir treatment, found in 6/8 hamsters. Scale bar, 100 µm. E. Peribronchial immune cell infiltration, mainly macrophages and lymphocytes, (green arrowhead), Nirmatrelvir treatment, found in 8/8 hamsters. Scale bar, 100 µm. F. Hyperplasia and hypertrophy of type II pneumocytes (green arrow), Nirmatrelvir-HA15 treatment, found in 7/7 hamsters. Scale bar, 100 µm.

